# Facilitating population genomics of non-model organisms through optimized experimental design for reduced representation sequencing

**DOI:** 10.1101/2021.03.30.437642

**Authors:** Henrik Christiansen, Franz M. Heindler, Bart Hellemans, Quentin Jossart, Francesca Pasotti, Henri Robert, Marie Verheye, Bruno Danis, Marc Kochzius, Frederik Leliaert, Camille Moreau, Tasnim Patel, Anton P. Van de Putte, Ann Vanreusel, Filip A. M. Volckaert, Isa Schön

**Author notes:** Correspondence: Henrik Christiansen.

## Abstract

Genome-wide data are invaluable to characterize differentiation and adaptation of natural populations. Reduced representation sequencing (RRS) subsamples a genome repeatedly across many individuals. However, RRS requires careful optimization and fine-tuning to deliver high marker density while being cost-efficient. The number of genomic fragments created through restriction enzyme digestion and the sequencing library setup must match to achieve sufficient sequencing coverage per locus. Here, we present a workflow based on published information and computational and experimental procedures to investigate and streamline the applicability of RRS. In an iterative process genome size estimates, restriction enzymes and size selection windows were tested and scaled in six classes of Antarctic animals (Ostracoda, Malacostraca, Bivalvia, Asteroidea, Actinopterygii, Aves). Achieving high marker density would be expensive in amphipods, the malacostracan target taxon, due to the large genome size. We propose alternative approaches such as mitogenome or target capture sequencing for this group. Pilot libraries were sequenced for all other target taxa. Ostracods, bivalves, sea stars, and fish showed overall good coverage and marker numbers for downstream population genomic analyses. In contrast, the bird test library produced low coverage and few polymorphic loci, likely due to degraded DNA. Prior testing and optimization are important to identify which groups are amenable for RRS and where alternative methods may currently offer better cost-benefit ratios. The steps outlined here are easy to follow for other non-model taxa with little genomic resources, thus stimulating efficient resource use for the many pressing research questions in molecular ecology.

## 1. Background

Evolutionary and ecological population genetic studies are important to understand how the diversity of life on earth is distributed, has evolved and may respond to future environmental changes (1). A grand challenge has been to document this biodiversity and understand its role in maintaining ecosystem functionality, particularly in the ocean (2) and even more so in frontier areas such as the deep-sea and polar regions (3). Molecular data collection has benefitted from a revolution in sequencing technologies such that genomics, where billions of nucleotides are screened simultaneously, is now an integral part of the biological toolbox (4–6). Genome-wide data open new avenues of ecological and evolutionary research, especially to study local adaptation (7,8). Given ever-increasing rates of anthropogenic disturbance, it is crucial to assess spatio-temporal genomic diversity, adaptation patterns and resilience of non-model organisms (9,10).

Similar to previous methodology shifts in population genetics (e.g. from Amplified Fragment Length Polymorphisms [AFLP] to microsatellites), the transition to novel methods requires detailed understanding of the new technology, its potential as well as its pitfalls, and careful experimental planning. While some study systems are moving towards population-specific shallow re-sequencing of whole genomes (e.g. important commercial fish species) (11,12), many species of interest with less extensive genomic resources rely on reduced representation sequencing (RRS) techniques to subsample the genome. These methods are attractive because they make more frugal use of sequencing volume. One or several restriction endonuclease enzymes are commonly used in RRS to first fragment the target genome into smaller portions, thus reducing sequencing costs. Millions of reads from high-throughput sequencing platforms are then aligned against either a reference genome or, alternatively, a *de novo* reference catalog of loci. Subsequently, genetic variants, most commonly single nucleotide polymorphisms (SNPs), but also microsatellites, introns, copy number variants (CNV) and microhaplotypes are determined (13–16).

In ecological and evolutionary research of natural populations, Restriction site-Associated DNA sequencing (RADseq) and derivatives (17–19) and Genotyping by Sequencing (GBS) (20,21) are the most commonly used terms for a variety of methods that essentially employ the same principle. The scientific nomenclature for such approaches lacks consistency (22,23). Here, we follow the reasoning of Campbell et al. (22) and use the term RRS (24) to refer to all of these methods. RRS has provided many important insights across a wide range of taxa from different ecosystems, e.g. with respect to population structure and demography, as well as hybridization, landscape or seascape genomics, QTL mapping, phylogeography, and shallow phylogenies (e.g. 5,25–30).

Effective and cost-efficient RRS experiments must be well designed. First, one should establish whether the species of interest truly represents one species or if cryptic species are present. This can be problematic in non-model taxa and has potentially large downstream implications for RRS such as high divergence but few shared loci (31,32). A useful complement is therefore DNA barcoding to screen for cryptic species (33,34). Alternatively, RRS can be specifically employed for species delimitation purposes (28,35,36), but this should be a deliberate choice before designing the RRS setup. For such a scenario it would be especially important to sequence many fragments thereby increasing the likelihood of capturing genetic markers that are conserved across, yet discriminatory between species. In general, the research question fundamentally determines whether the application of RRS is appropriate. For example, providing evidence for significant, evolutionary neutral genetic population structure may be easier and less expensive with a good number (>10) of multi-allelic microsatellites (37). However, RRS may be better suited to identify loci that are putatively affected by spatially variable selection and therefore involved in local adaptation. To this end, the density of markers (SNPs across the genome) that can be realized for a given species, which depends on genome size and complexity, as well as research budget, should be considered.

With low marker density one may run the risk of accepting unreasonably high rates of false positives (outliers that are not based on biological reality) in genome scans leading to biased or erroneous inferences (38,39). Consequently, there is debate about the usefulness of RRS (or RADseq in particular), especially for inferring local adaptation patterns (40,41). The genomic characteristics of a target species, most importantly its genome size and the level of linkage disequilibrium (LD), are crucial to design a RRS experiment. With little genomic information, *a priori* calculations may be inaccurate. Therefore, it is vital to assess, optimize and critically ponder the advantages and limitations of RRS for a given research project to avoid the creation of sequence data that are unsuitable to answer the study question and/or inefficient use of resources. A most critical point is to properly strike a balance between sequencing depth (coverage) and number of fragments, which is roughly proportional to the number of genetic markers. The estimated number of fragments generated from a genome determines the marker density (as the number of fragments translates approximately into the number of SNPs), while avoiding unnecessary “over”-sequencing of the genomic fragments, i.e. loci or RADtags, to save sequencing costs. Both excessive (>100×) and uneven or too low (<10×) coverage is detrimental for accurate locus reconstruction and SNP calling, particularly in *de novo* approaches (42). Hence, RRS experimental procedures may benefit from thorough optimization. In this context, we used the framework of a large research project (“Refugia and Ecosystem Tolerance in the Southern Ocean”) to optimize RRS for a diverse set of taxa in parallel. The Southern Ocean hosts a unique marine fauna with high levels of endemism (43,44), but is increasingly subject to external pressures, such as warming, pollution and living resource exploitation (45–48). Population genomic approaches are needed to understand the genetic structure and connectivity of Antarctic fauna, so that appropriate management and conservation actions can be developed (e.g. 49–51).

In this molecular pilot experiment, we seek to investigate and optimize the applicability of RRS to a range of Antarctic non-model taxa across the animal kingdom. The target organisms are ecologically important, abundant, and widely distributed in the Southern Ocean and cover a variety of habitats – from benthos to pelagic birds. Specifically, we aim to develop economic and robust experimental setups for RRS population genomic studies in an ostracod group (*Macroscapha opaca-tensa* species complex, Brandão et al., 2010) (Ostracoda, Macrocyprididae), two amphipod species (*Charcotia obesa* Chevreux, 1906 and *Eusirus pontomedon* Verheye & D’Udekem D’Acoz, 2020) (Malacostraca, Lysianassidae and Eusiridae), two bivalve species (*Laternula elliptica* P.P. King, 1832 and *Aequiyoldia eightsii* Jay, 1839) (Bivalvia, Laternulidae and Sareptidae), two sea star species (*Bathybiaster loripes* Sladen, 1889 and *Psilaster charcoti* Koehler, 1906) (Asteroidea, Astropectinidae), two fish species (*Trematomus bernacchii* Boulenger, 1902 and *Trematomus loennbergii* Regan, 1913) (Actinopterygii, Nototheniidae), and the two snow petrel subspecies (*Pagodroma nivea nivea* Forster, 1777 and *Pagodroma nivea confusa* Clancey, Brooke & Sinclair, 1981) (Aves, Procellariidae). The outlined approach should be readily adoptable for other taxa of interest. We lay out a clear and concise protocol to follow *a priori* for any RRS experiment on non-model species that will help researchers to evaluate the costs, benefits, and risks of such projects.

We specifically aim to (i) collate information about the genomic properties of the target taxa; (ii) assess *in silico* which restriction enzymes are likely to yield the desired number of fragments; (iii) test selected restriction enzyme digestions in the laboratory; (iv) optimize restriction enzyme choice, size selection window and the number of individuals to be pooled per sequencing library (based on the previous results); and (v) sequence and analyze test RRS libraries of promising experimental setups. These extensive pilot analyses – including literature research, computational analyses, and laboratory work – are designed to comprehensively evaluate all information for each target species or species complex. In the workflow of optimizing the setup for each target taxon, we strive to use the same restriction enzymes (or combinations) for several taxa whenever possible to reduce the costs for specifically designed barcodes and adaptors. Results shall ultimately facilitate informed decisions about whether and how RRS for each taxon could be conducted. We critically discuss these considerations and suggest alternative approaches in two cases.

## 2. Methods

### 2.1 Specimen sampling

Samples of all target species were available from recent expeditions to the Southern Ocean (Additional File 1). For ostracods, we used existing DNA extractions of Macrocyprididae from the Southern Ocean that were already taxonomically identified and described (52,122). The amphipod target species were collected during RV *Polarstern* (123) expedition ANTXXIX-3 PS81. More details on *Eusirus pontomedon* (note that we initially included these specimens tentatively as *Eusirus* aff. *perdentatus*, but the taxonomy was updated during the course of this project) are provided in (124), while details of investigated *Charcotia obesa* are given in (125). The bivalves *Laternula elliptica* and *Aequiyoldia eightsii* were sampled by scuba diving in the shallow water of Potter Cove (King George Island, western Antarctic Peninsula; by F. Pasotti) and Rothera station (Adelaide Island, West Antarctic Peninsula; courtesy of the British Antarctic Survey) in 2016. Two sea star species (*Bathybiaster loripes* and *Psilaster charcoti*) were collected during international expeditions with RRS *James Clark Ross* and RV *Polarstern* to the South Orkney Islands (JR15005 in 2016, PS77 in 2011), the Weddell Sea (PS81 in 2013), West Antarctic Peninsula (PS77 in 2011), and with RV *L’Astrolabe* to Adélie Land (REVOLTA 1 in 2010). Emerald rockcods (*Trematomus bernacchii*) were sampled in 2014 around James Ross Island with gill nets (126). Scaly rockcods (*Trematomus loennbergii*) were sampled in the Ross Sea as bycatch of the exploratory Antarctic toothfish (*Dissostichus mawsoni*) longline fishery. Dead birds and feathers of snow petrels (*Pagodroma* spp.) were sampled during the BELARE 2017-2018 expedition in the vicinity of the Princess Elisabeth Station, and additional samples were obtained from Signy and Adelaide Islands as courtesy of the British Antarctic Survey. Samples were stored frozen, dried, or in >90% ethanol until DNA extraction.

### 2.2 Genomic resources

Prior to computational analyses, genomic information was collated for all target species or, if such information was not available, from the closest related species. Published reference genomes were collected from the literature and online resources, such as GenBank and Ensembl (127). In addition, genome size estimates were retrieved from genomesize.org (128) and other published estimates based on flow cytometry (e.g. 67). Genome size estimates as C values were transformed to Mb for comparison (1 pg = 978 Mb) (95).

### 2.3 *In silico* genome digestion analyses

We used SimRAD to computationally digest genomic DNA at sites matching a restriction enzyme recognition site (130). In total, seven restriction enzymes and combinations thereof were tested (Table 1). These were chosen based on what is commonly used in comparable studies and to cover a variety of enzymes ranging from very common (*MseI, MspI, ApeKI*) to medium (*EcoRI, SphI, PstI*) and rare cutters (*SbfI*). Reference genomes from related species as well as two simulated genomes per taxonomic class were used for these *in silico* digestions. Simulated genomes were modelled with GC content as in the available reference genome(s) and with two different sizes per taxonomic class to cover the approximate range of genome sizes known for this class (Table 2). The total number of fragments that these enzymes (or enzyme combinations for double digest setups) produced were estimated, as well as the number of fragments in various size selection windows (between 210-260, 240-340, 0-100, 100-200, 200-300, 300-400, 400-500, 500-600, 600-700, 700-800, and 800-900 bp). Approximate targets for the number of fragments in each species of interest were defined (Table 2) and restriction enzyme and size selection combinations that fell within the target range ± 10,000 fragments were retained for downstream testing. After narrowing down the enzyme choice and conducting empirical digestion analyses, we ran additional *in silico* digestions for a final optimization of the size window and thus number of fragments for each specific case. During these fine-tuning analyses we tested as many different size selection windows as needed (in some cases >20 additional size windows between 50 and 250 bp width) to find a suitable estimate of the number of fragments.

**Table 1.** Genomic information useful for reduced representation sequencing (RRS) optimization in target species from six organism classes.

| Class | Target Species | Target Fragment Number | Genome Size Estimates (C) | Available Genome Size (Mb) from related species | Simulated Genomes |
| --- | --- | --- | --- | --- | --- |
| *Ostracoda* | Macrocyprididae | 50,000 | 0.17±0.003<sup>†</sup> | *Cyprideis torosa*, 286 Mb (54) | 100 Mb, 43.9 % GC<br>500 Mb, 43.9 % GC |
| *Malacostraca* | *Charcotia obesa*<br>and *Eusirus pontomedon* | 50,000 | unknown<br>(Amphipoda: 0.68-64.62<sup>†</sup>) | *Hyalella Azteca*, 551 Mb (56)<br>*Parhyale hawaiiensis*, 4,003 Mb (57) | 10,000 Mb, 38.5 % GC<br>30,000 Mb, 40.8 % GC |
| *Bivalvia* | *Laternula elliptica*<br>and *Aequiyoldia eightsii* | 50,000 | unknown<br>(0.65-5.40<sup>†</sup>) | *Crassostrea gigas*, 558 Mb (60)<br>*Pinctada imbricata*, 991 Mb (61)<br>*Bathymodiolus platifrons*, 1,658 Mb (62) | 1,000 Mb, 35.3 % GC<br>5,000 Mb, 34.2 % GC |
| *Asteroidea* | *Bathybiaster loripes* and *Psilaster charcoti* | 20,000 | unknown<br>(Asteroidea: 0.54-0.96<sup>†</sup>) | *Acanthaster planci*, 383 Mb (63)<br>*Patiria miniata*, 811 Mb (64)<br>*Patiriella regularis*, 949 Mb (65) | 1,000 Mb, 41.3 % GC<br>2,000 Mb, 40.4 % GC |
| *Actinopterygii* | *Trematomus bernacchii* and *T. loennbergii* | 20,000 | *T. bernacchii*: 1.12±0.019<sup>‡</sup>; 1.19<sup>§</sup>; 1.82<sup>¶</sup>; *T.* | *Notothenia coriiceps*, 637 Mb (58) | 1,000 Mb, 40.8 % GC<br>1,800 Mb, 40.8 % GC |
|  |  |  | *loennbergii*:<br>1.34<sup>‡</sup> |  |  |
| Aves | *Pagodroma nivea*<br>*nivea* and *P. nivea*<br>*confusa* | 50,000 | unknown<br>(0.91-2.16<sup>†</sup>) | *Fulmarus glacialis*, 1,141 Mb<br>(59) | 1,500 Mb, 41.2 % GC<br>2,000 Mb, 41.2 % GC |

For each class approximate targets for the number of fragments were defined and known genome size estimates from flow cytometry are listed. In species with unknown genome size, the range of published estimates from species from the same class is listed. Available genomes from related species and two simulated genomes per class were used for *in silico* digestions. The simulated genomes were modeled according to expectations regarding genome size and GC content as known from related species.
<sup>†</sup> published estimates from various species of the same class (or where indicated order), as listed on genomesize.com on 9<sup>th</sup> January 2019; Ostracoda: Macrocyprididae: Jeffery et al., 2017
<sup>‡</sup> Auvinet et al., 2018
<sup>§</sup> Hardie and Hebert, 2003
<sup>¶</sup> Morescalchi et al., 1996

**Table 2.** Restriction enzymes and combinations used for reduced representation sequencing (RRS) optimization.

| Restriction Enzyme<br>(Combination) | Recognition Site | Approximate<br>Fragment Number<sup>†</sup> | Special Features | Reference |
| --- | --- | --- | --- | --- |
| *SbfI* | 5'--CCTGCA\|GG--3' | 6,000 |  | e.g. (35,69–71) |
| *EcoRI* | 5'--G\|AATTC--3' | 320,000 | Methylation<br>sensitive | (19,72) |
| *SphI* | 5'--GCATG\|C--3' | 150,000 |  |  |
| *PstI* | 5'--CTGCA\|G--3' | 150,000 |  | (73,74) |
| *ApeKI* | 5'--G\|CWGC--3' | 1,000,000 | Methylation<br>sensitive, degenerate site | e.g. (75–78) |
| *MspI* | 5'--C\|CGG--3' | 1,700,000 |  |  |
| *MseI* | 5'--T\|TAA--3' | 7,600,000 |  | (79) |
| *SbfI_SphI* |  | 12,000 |  | e.g. (80–83) |
| *SbfI_MspI* |  | 12,000 |  | (84,85) |
| *PstI_MspI* |  | 275,000 |  | e.g. (21,86–88) |
| *EcoRI_SphI* |  | 250,000 |  | (89) |
| *EcoRI_MspI* |  | 540,000 |  | e.g. (90–93) |

Recognition site, the approximate expected fragment number in a 1000 Mb genome, any special enzyme characteristic and empirical studies that recently used this enzyme (combination) are listed.
† for a 1000 Mb genome with 40 % GC content and no size selection

### 2.4 Empirical genome digestion analyses

Laboratory experiments were conducted with promising restriction enzymes to complement results from computational analyses. For each species, DNA from three individuals was used to test two or three restriction enzymes or enzyme combinations. Genomic DNA was extracted using either the commercial DNA extraction kits NucleoSpin Tissue (Macherey-Nagel) or DNeasy Blood & Tissue (Qiagen) and following the manufacturer’s guidelines, or with a standard salting out protocol (81), or, for the bivalves, with a standard cetyl trimethylammonium bromide (CTAB) protocol. Subsequently, DNA quality and quantity were checked using the fluorescence assay Quant-iT PicoGreen dsDNA (Thermo Fisher Scientific Inc.), an Infinite M200 microplate reader (Tecan Group Ltd.) and 1 % agarose gel electrophoresis. Whenever possible, only high-quality DNA extractions were used. Because of their small size, extractions from individual ostracods yielded insufficient quantities of DNA for downstream protocols, and sample numbers per locality were very low. Hence, the entire genomic DNA of ostracods was amplified using the REPLI-G kit (Qiagen) for whole genome amplification of 1 μL extracted DNA with high-fidelity polymerase Phi 20 and multiple displacement amplification following the manufacturer’s protocol. For this purpose, extractions with the highest DNA concentrations from different species of Macrocyprididae, mainly of the *Macroscapha tensa-opaca* species complex, were selected (52). For all target species, 100 ng genomic DNA of three biological replicates per species was digested with 10 units of a selected restriction enzyme at 37 °C (*EcoRI, MspI* and *PstI*) or 75 °C (*ApeKI*) for 2 h in a total volume of 10 μL. Reactions were purified with CleanPCR (GC Biotech) according to the manufacturer’s protocol. Between 1 and 5 ng of the purified digested DNA was loaded on a High Sensitivity DNA chip (Agilent Technologies) and run on an Agilent 2100 Bioanalyzer System. Results were exported from the 2100 expert software (Agilent) as XML files and read into R v4.0.4 (131) using the bioanalyzeR package v0.5.1 (132). Additional R packages used in this project were here v1.0.1 (133), seqinR v1.0-2 (134), the tidyverse packages (135), ggsci v2.9 (136), and gridExtra v2.3 (137) (see also more details under: https://github.com/notothen/radpilot). Because it is not possible to accurately standardize the number of fragments in an empirical digest without knowledge of the true genome size, we compared the shape of the curves of produced fragments (number of loci or DNA concentration vs. locus size or length) between *in silico* and empirical digests (Fig. 2 and Additional File 3).

**Figure 2.**
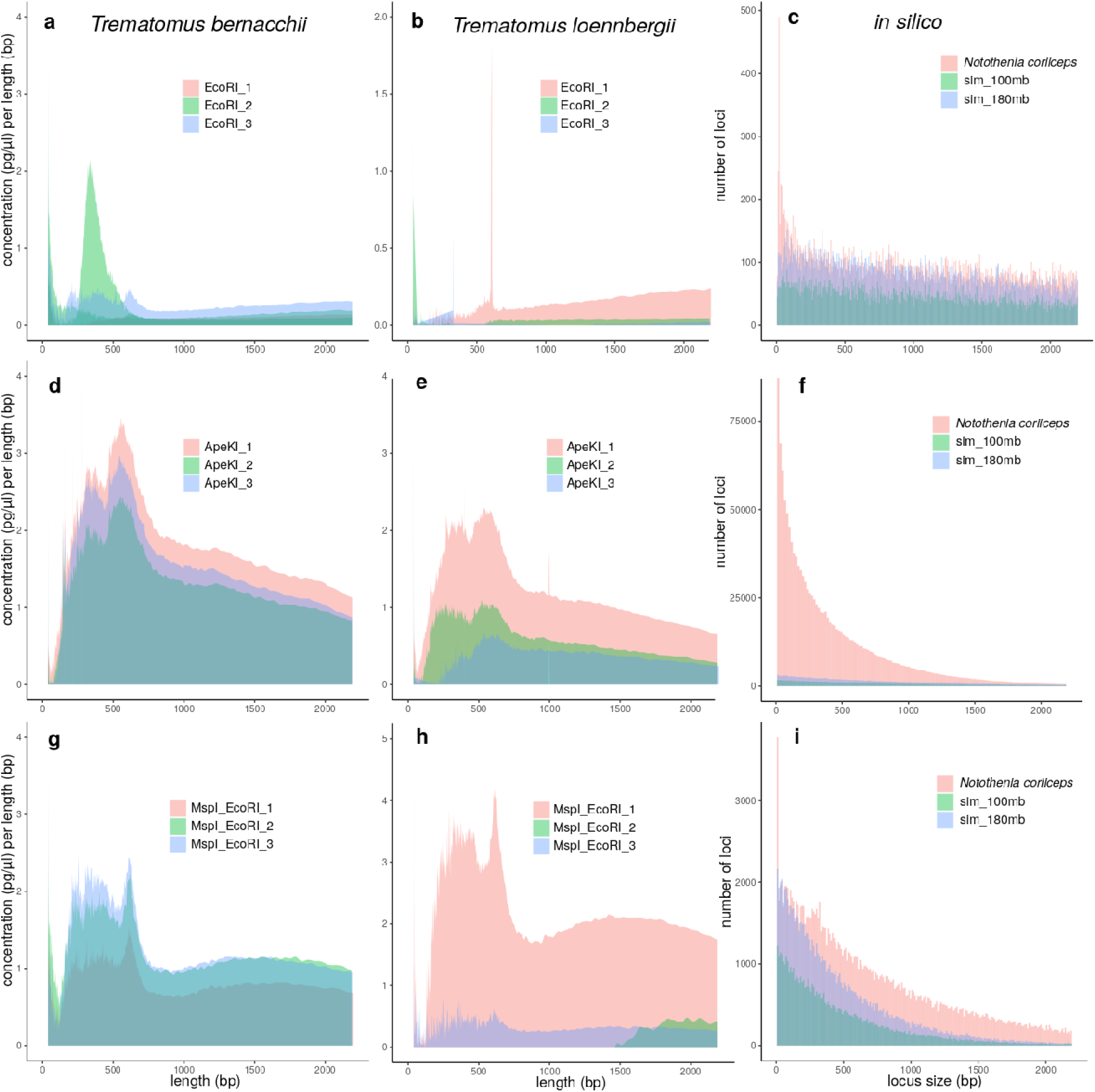
Comparison of empirical and *in silico* restriction enzyme digestions. Empirical Bioanalyzer results with digested DNA are shown as concentration over fragment size (a, b, d, e, g, h) and estimated loci numbers over locus size from *in silico* digestions (c, f, i). The tests were conducted with restrictions enzymes *EcoRI* (a, b, c), *ApeKI* (d, e, f) and a double digest with *EcoRI* and *MspI* (g, h, i). Results for the fish species *Trematomus bernacchii* (a, d, g) and *T. loennbergii* (b, e, h) are shown next to *in silico* estimates using a related reference genome of *Notothenia coriiceps* and two simulated genomes of 100 and 180 Mb size (note that this was the absolute size used for *in silico* computations, but resulting estimates were extrapolated to 1000 and 1800 Mb). Results for other target taxa are shown in Additional File 3.

### 2.5 RRS setup optimization

In order to choose a promising restriction enzyme and size selection combination, we calculated the sequencing coverage per fragment as follows:

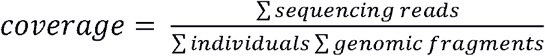

We conservatively aimed at a coverage of approximately 30× for each fragment per individual, higher than other minimum recommendations (17,32,42). Given that the accuracy of our genome size estimates is unknown, we aimed for relatively high coverage, so that in a “worst-case scenario”, where the genome size is actually twice as large as we estimated (or any other factor leads to twice as many fragments as assumed), we would still reach a coverage of approximately 15×. The number of individuals per sequencing library was set to 96, corresponding to one PCR plate. Sequencing with a HiSeq 4000 platform (Illumina) should conservatively yield approximately 300 million reads per sequencing lane, while on a HiSeq 2500, we expect approximately 200 million reads. These coverage calculations were applied to fragment numbers from *in silico* results based on available reference genomes and extrapolated to a final, conservative estimate of genome size based on the best available knowledge (Table 3). This extrapolation is likely not biologically accurate but serves as a conservative correction factor. We then used *in silico* estimates again to further tweak the size window of a chosen restriction enzyme or enzyme combination in each target species to achieve the desired coverage, while considering the size range in which the two HiSeq machines work best. Finally, we estimated the number of SNPs across the genome as a measure of marker density (analogous to 40) for a chosen enzyme and size selection setup and sequencing machine, assuming one SNP every other 100 bp. The latter estimate is based on our own experience, predominantly from fish genomes (but see also e.g. 99). Spreadsheet tables and R scripts for the main calculations, data and plots regarding *in silico* analyses, coverage and marker density are available at https://doi.org/10.5281/zenodo.3267164 and at https://github.com/notothen/radpilot.

**Table 3.**
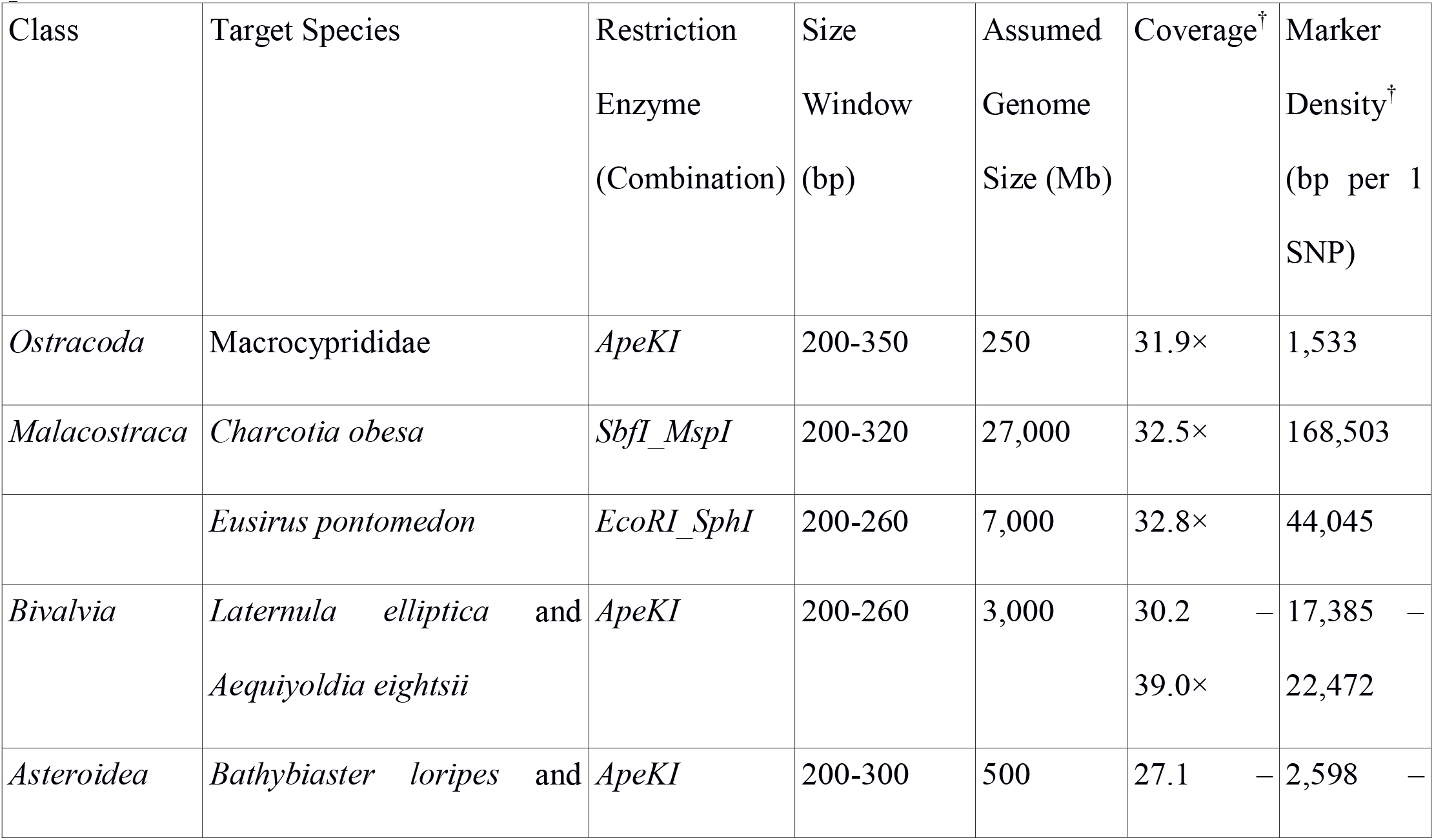
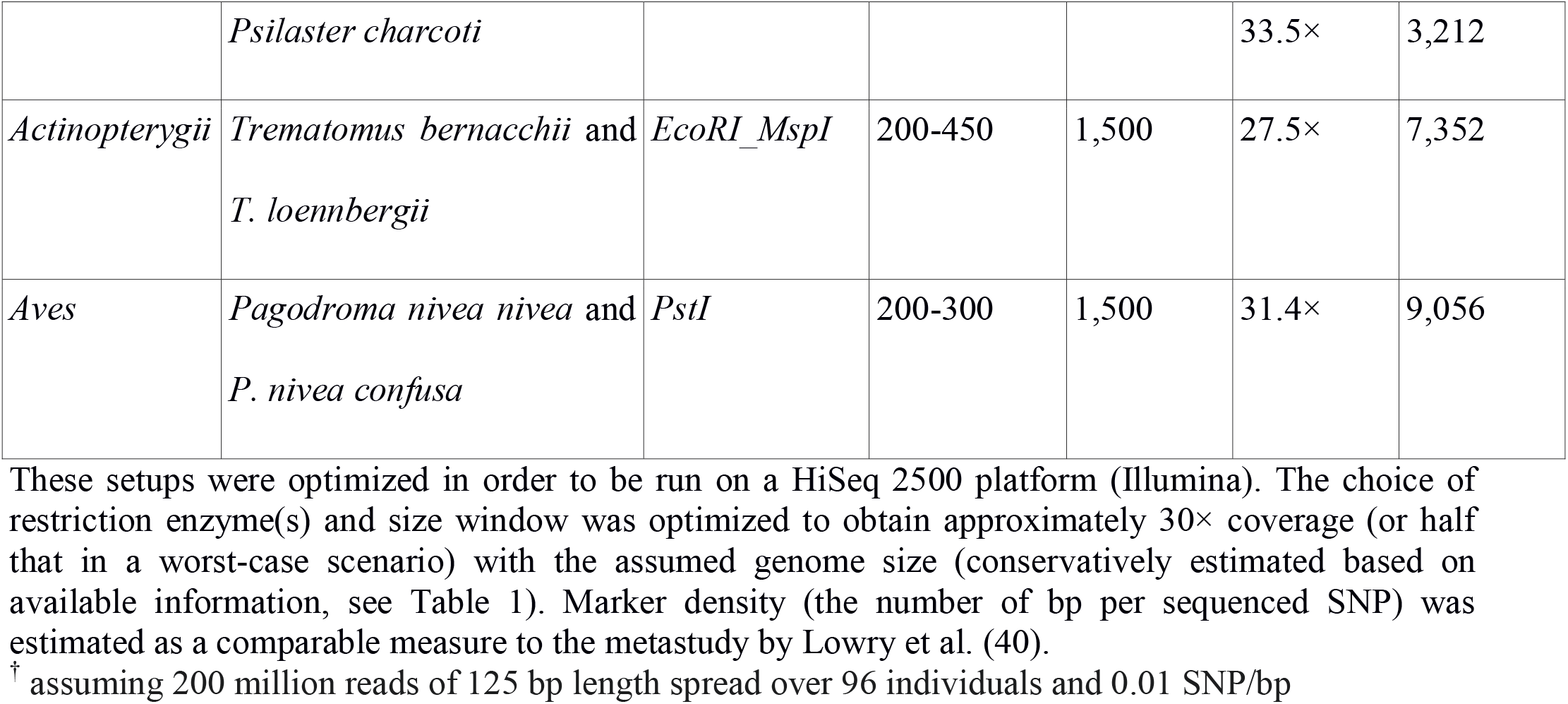
Reduced representation sequencing (RRS) setups for seven individually optimized protocols.

### 2.6 RRS library preparation and sequencing

The information collected so far convinced us not to pursue RRS in amphipods (see discussion); they were therefore not included in the test libraries. In addition, not enough high molecular weight DNA samples of *P. nivea confusa* (one of the snow petrel subspecies) were available. Eventually, five RRS test libraries for eight target species were constructed using 6, 8, 10, or 14 individuals and two controls per species and sequenced on one lane of a HiSeq 2500 unit (see Table 4 in results section). That way, we attempted to keep the fixed variables for our coverage calculations, i.e. an estimated 250 million reads spread over 94 individuals and between 53,399 and 81,605 fragments. We originally aimed at 96 individuals, but too many samples of low-quality DNA dropped out during sample preparation. In addition, the estimated number of fragments varied between target species, but the conservative estimates in all other aspects should allow for some flexibility here. The libraries were all prepared by the same person at the KU Leuven laboratory using custom protocols that are based on two main references: the original ddRAD protocol by Peterson et al. (17) and the original GBS protocol by Elshire et al. (20). We adjusted these protocols slightly and provide a full-length description of the laboratory procedure in Additional File 5 & 6. In both cases, the standardized high-quality DNA was first digested with restriction enzyme(s), followed by adaptor and barcode ligation, purification, PCR, another purification and finally quantification and pooling. The libraries were then sent to the KU Leuven Genomics Core (www.genomicscore.be), where all five libraries were individually size selected on a Pippin Prep unit (Sage Science), checked for quantity using qPCR, pooled, and paired-end sequenced on one lane of a HiSeq 2500 platform (Illumina).

**Table 4.** Setup and results of five test libraries for reduced representation sequencing (RRS) from eight species/groups.

| Library Nr. | Class | Target Species | Protocol, Enzyme and Size Window (bp) | N + controls | Stacks parameter *M* and *n* | Expected Nr. of Fragments<sup>†</sup> | Obtained Loci<sup>†</sup> | Expected Coverage | Obtained Coverage<sup>†</sup> |
| --- | --- | --- | --- | --- | --- | --- | --- | --- | --- |
| 1 | *Ostracoda* | Macrocyprididae | GBS, *ApeKI*, 200-350 | 8 + 2 | 6 | 65,244 | 69,817<br>(±63,114) | 31.9× | 28.2×<br>(±5.4) |
| 2 | *Bivalvia* | *Laternula elliptica*<br>*Aequiyoldia eightsii* | GBS, *ApeKI*, 200-260 | 8 + 2<br>10 + 2 | 4 | 53,399 – 69,027 | 125,305<br>(±22,828)<br>143,551<br>(±28,676) | 30.2 – 39.0× | 21.6×<br>(±5.1)<br>20.0×<br>(±2.6) |
| 3 | *Asteroidea* | *Bathybiaster loripes* | GBS, *ApeKI*, 200-300 | 10 + 2<br>14 + 2 | 5 | 62,272 – 76,988 | 82,945<br>(±43,521) | 27.1 – 33.5× | 21.0×<br>(±6.8) |
|  |  | *Psilaster*<br>*charcoti* |  |  |  |  | 115,608<br>(±30,589) |  | 27.6×<br>(±4.6) |
| 4 | *Actinopterygii* | *Trematomus*<br>*bernacchii*<br>*T. loennbergii* | ddRAD,<br>*EcoRI_MspI*,<br>200-450 | 10 + 2<br>10 + 2 | 3 | 81,605 | 21,121<br>(±3,539)<br>23,609<br>(±2,362) | 27.5× | 42.3×<br>(±13.5)<br>49.6×<br>(±6.2) |
| 5 | *Aves* | *Pagodroma*<br>*nivea nivea* | GBS, *PstI*,<br>200-300 | 6 + 2 | 3 | 66,258 | 140,972<br>(±26,444) | 31.4× | 10.0×<br>(±0.4) |

‡ only estimates from real (not from simulated) genomes listed
† using denovo\_map.pl with $m=3$ and $M=n$ as listed in column six

### 2.7 Sequence analyses

Sequencing data were checked using FastQC v0.11.5 (139) and then demultiplexed and cleaned (options -c and -q) using the process_radtags module of Stacks v2.4 (95,96). Because some of our multiplexing barcodes for the *PstI* library were contained in longer *ApeKI* barcodes, we demultiplexed the *ApeKI* libraries first and captured reads that were discarded in the process. These reads were subsequently used for demultiplexing of the *PstI* library. All demultiplexing runs were conducted without barcode rescue to avoid cross-contamination between libraries. The Stacks pipeline was also used for each target species independently to create a *de novo* assembly and call genotypes. Building contigs from paired-end reads is not possible with GBS data in Stacks (15), because the orientation of the reads is ambiguous. In this case (libraries 1, 2, 3, 5), we concatenated the four output files per individual of process_radtags to run the pipeline as if it was single-end data. Our size selection windows were designed to avoid overlap between the two reads of one fragment, so this approach should work well, albeit creating shorter haplotypes. We used Stacks’ default value for *m*, i.e., a minimum stack coverage of 3, which generally produces consistent results at typical coverage rates (32). Choosing parameters *M* and *n* to control the formation of loci within and across individuals on the other hand is study dependent. We explored a parameter range of *n*=*M*=[1 .. 9] following Rochette and Catchen (42) to strike a balance between over- and undermerging alleles and loci. To compare results from the different parameters only loci present in 80 % of the samples (50 % in the case of ostracods) were retained. Further detailed filtering would be required for downstream population genomic analyses.

## 3. Results

The optimization process of RRS experimental setups for non-model species is iterative and includes many deliberate choices that must be made based on the best available knowledge (Fig. 1). Relatively constant variables, i.e. the number and quality of samples, the research budget and the main research question, should be considered during the entire process and flexible variables, such as restriction enzymes, size selection window and the number of individuals to be pooled, should be adjusted to reach the desired outcome.

**Figure 1.**
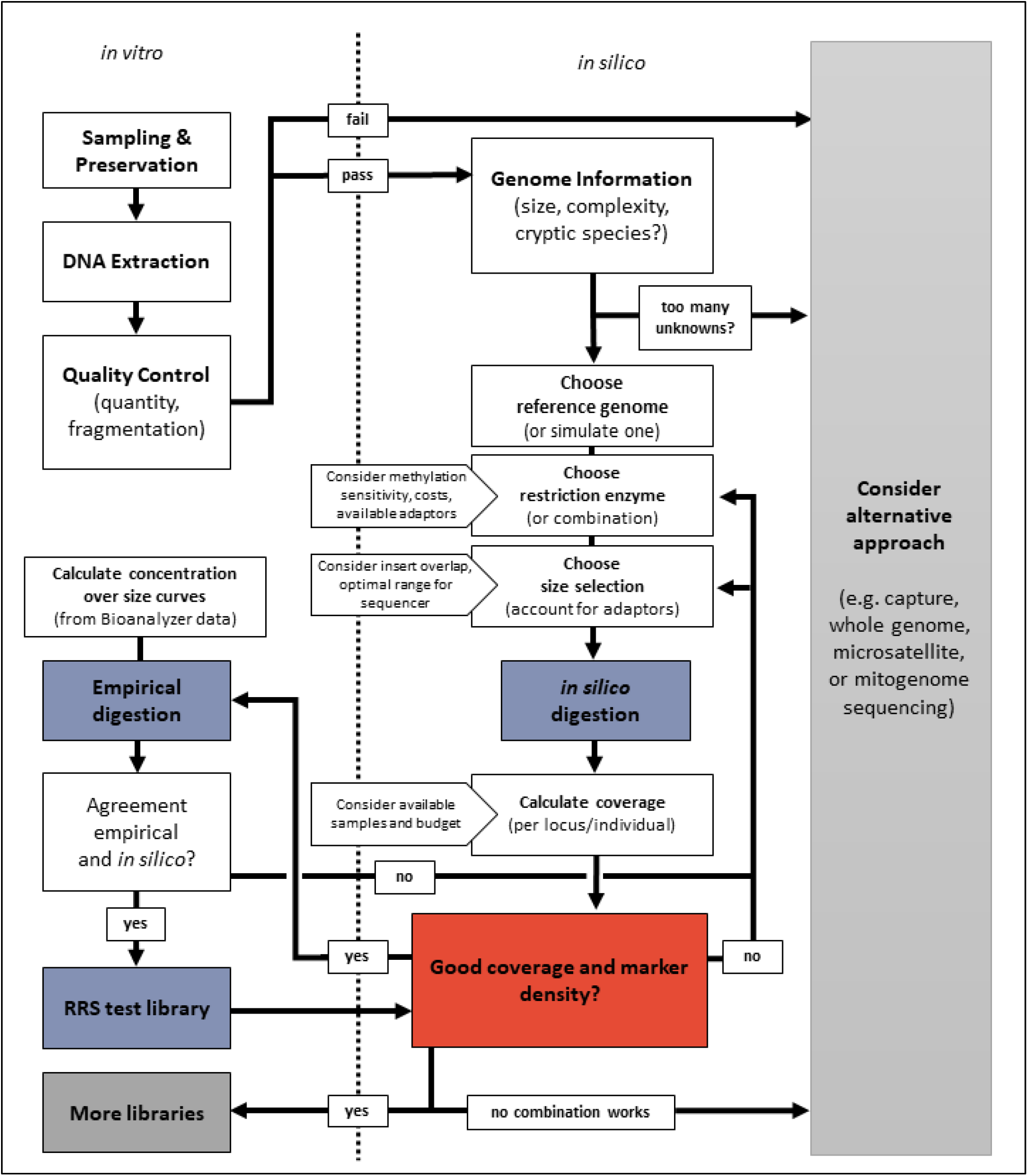
The iterative process of reduced representation sequencing (RRS) optimization. Empirical (*in vitro*, left of the dotted line) and computational (*in silico*, right of dotted line) analyses are part of this process. Core procedures to identify suitable experimental setups are *in silico* and empirical enzyme digestion and sequencing of a pilot RRS library (blue boxes). The coverage and marker density that can be achieved with a given setup needs to be repeatedly checked and fine-tuned (red box).

### 3.1 Genome characteristics

First, published genomic resources of our target taxa were collected. Available information is highly variable across target taxa, with typically more genomic resources available for vertebrate groups (Table 1). Genome size among ostracods varies considerably, with Macrocyprididae estimated at approximately 166 Mb (or 0.17 C) (53). One published ostracod genome (*Cyprideis torosa*) with a genome size comparable to our target species was available (54). Amphipods show very large variability in genome size (55) with extreme cases that dramatically exceed the size estimates of all other target taxa studied here (up to 63,198 Mb or 64.62 C, Table 1). Two amphipod reference genomes were available (*Hyalella azteca, Parhyale hawaiensis*) (56,57). For bivalves and sea stars more reference genomes were available, but not from species closely related to the target species. In both cases, we selected three reference genomes of varying size (Table 1). The Antarctic fish target species have genome size estimates available as well as a reference genome from a species from the same family (*Notothenia coriiceps*) (58). In birds, no genome size estimates for our target species were available, but bird genome size appears to be relatively constrained between approximately 1 and 2 Gb and a reference genome from the same family has been published (*Fulmarus glacialis*) (59). We decided to aim at 50,000 fragments as initial targets for our optimizations in all taxa, except for fish and sea stars (Table 1). In the latter target taxa, we aimed at 20,000 fragments initially, because we had more samples available and thus were interested in covering more individual samples from a wider geographic range at the expense of marker density. Note that these targets are highly study specific and depend on the budget, number of samples to be sequenced and, most importantly, exact research question of a given RRS project.

### 3.2 *In silico* digestions

We estimated how many RRS fragments twelve restriction enzymes and enzyme combinations (listed in Table 2) would produce. These estimates were conducted using various reference genomes and simulated genomes. We estimated the fragment number in total as well as in various size selection windows (see below and Additional File 2). As expected, the fragment number is influenced primarily by the type of enzyme and the genome size. The tested combinations produced fragment numbers close to our defined targets in all species, but with different restriction enzymes. We aimed at using as few different enzyme setups across species as possible. Using the same setup for several RRS experiments reduces costs as the same adaptor sets can be reused multiple times. Therefore, we kept five initial setups that yielded promising fragment numbers: *EcoRI, PstI, ApeKI, MspI* and a double digest with *EcoRI* and *MspI*.

### 3.3 Empirical digestions

Based on preliminary *in silico* results, we tested the genome digestion by several enzymes and enzyme combinations in the laboratory. High quality bird DNA was not available, preventing empirical digestion tests for this group. Ostracod DNA was whole genome amplified and this proved problematic for the Bioanalyzer instrument, because the results indicated overloading even after multiple dilutions. In total, 75 empirical digestions were conducted, several of which produced unusable results even after repeating the experiment. The sensitivity of the Bioanalyzer to small irregularities especially in the size range below 500 bp made it impossible to infer sensible patterns in many cases (Additional File 3). We therefore must caution that Bioanalyzer results only sometimes provide useful additional information that increase the confidence in estimates obtained *in silico*. Nevertheless, from the successful runs it appeared that the empirical results are usually more similar to *in silico* digestions with genomes from related species than of simulated genomes (Fig. 2 and Additional File 3). For example, in fishes *ApeKI* was estimated to produce significantly more small than large fragments using the *N. coriiceps* reference genome, which was at least roughly confirmed through empirical digestion (Fig. 2). Here, using *EcoRI* provides few fragments overall, which proved difficult to accurately depict using the Bioanalyzer. In contrast, the tested double digest provided a consistent picture in four out of six replicates for the two fish species (Fig. 2). Here, we also noted a pronounced spike at around 650 bp and the size window was therefore deliberately kept lower (see below and Table 3 and 4).

### 3.4 RRS setup

With all information gathered thus far, we proceeded to optimize the RRS experimental setup for each of the target taxa. We planned the same setup for species from the same class, when the genomic differences between those species were unknown (in Bivalvia and Asteroidea), or when they were related and therefore likely to have similar genomic properties (Actinopterygii and Aves). In contrast, we designed two different setups in Malacostraca, because the genomes of *C. obesa* and *E. pontomedon* may have very different sizes (Table 1 & 3). Experimental setups, i.e. restriction enzymes and size selection window, were furthermore tuned to suit a sequencing experiment with the HiSeq 2500 or 4000 platforms, respectively. The choice of sequencing platform can be modified based on instrument availability and budget. In the following, results for use with a HiSeq 2500 platform are listed (Table 3), the same results for a HiSeq 4000 platform can be found in Additional File 4. The setup for optimizing results as listed here also includes the consideration that it would be cost-efficient to use the same enzyme or enzyme combinations for several species whenever possible, because adaptors can then be re-used. Therefore, when several enzymes (or combinations) seemed promising according to *in silico* digestion, we attempted to choose setups that were also promising in other target species. For ostracods, we assumed a genome size of 250 Mb and 500 Mb as worst-case scenario. Using the *Cyprideis torosa* reference genome, a digest with *ApeKI* and size selection of 200 – 350 bp would yield 31.9× coverage (or half of that in the worst-case scenario). With this setup and genome size, we would achieve an estimated marker density of approximately one SNP every 1.5 kb. In amphipods, different setups per species are required. Given the highly uncertain genome size of 27,000 Mb for *C. obesa* and 7,000 Mb for *E. pontomedon* (based on same family estimates; 55), double digest RADseq experiments with *SbfI* and *MspI* and *EcoRI* and *SphI*, respectively, would yield the desired coverage. Marker density in both cases is expected to be low, due to the large genome size (Table 3). Because of uncertainty with respect to genome size and an anticipated low marker density, we stopped RRS optimization in amphipods and instead explored alternatives as envisioned in our workflow (Fig. 1). For both bivalve species, a genome digestion with *ApeKI* and size selection of 200 – 260 bp seemed promising with all three reference genomes and would yield around one SNP per 20 kb. Similarly, in sea stars we found setups with *ApeKI* and a slightly wider size selection that should yield good results, although results varied depending on the reference genome used. For the Antarctic fishes of the genus *Trematomus*, a double digest setup with *EcoRI* and *MspI* in a size window of 250 – 450 bp should yield desired coverage and marker density. Regarding the snow petrels, a setup with *PstI* and 200 – 300 bp size selection seemed appropriate, yielding one SNP every 9 kb. Overall, results indicate that with only three enzyme choices, it should be possible to achieve the desired coverage and marker density in eight species (Table 4).

### 3.5 RRS test libraries

Pilot libraries with optimized setups were sequenced, yielding a total of 531 million (M) reads. After demultiplexing and quality control, 422 M reads were retained. These reads were spread relatively evenly across libraries, species, and individuals (average and standard deviation across all taxa and libraries: 4.5 ± 2.1 M reads). All but five individuals received more than 1 M reads and most individuals received more than 3 M reads. We created *de novo* catalogs from these reads using Stacks (15,95,96) with varying *M* and *n* parameters (42). Optimal parameters varied (*M* = *n* = 3-6) among taxa (Table 4 & Additional File 7). Results from this parameter optimization also revealed varying levels of diversity, e.g. sea stars showed relatively high levels of polymorphism, while the bird library produced many loci but few SNPs (Additional File 7). Comparing the unfiltered numbers of loci and coverage across individuals underlined the inverse relation of these two variables (Table 4, Fig. 4). In ostracods, our target estimates were matched best. In bivalves and sea, stars more loci than expected were sequenced at the expense of coverage, although coverage was still reasonable. Two individuals of *B. loripes* had low coverage due to low initial numbers of reads, indicating errors during library preparation or degraded input DNA. The fish libraries contained considerably less loci than expected at high coverage, while the opposite was true for the bird library. The latter also showed very uniform low coverage at approximately 10×. Overall, these results show promise for full scale RRS libraries with sufficiently high coverage in four of five libraries.

## 4. Discussion

High-throughput sequencing methods promise new avenues of ecological and evolutionary research in non-model organisms. We provide a detailed workflow to evaluate and optimize reduced representation sequencing (RRS) techniques for any animal species of interest (Fig. 1). This approach is reproducible and ensures that researchers are well-informed about the advantages and drawbacks of RRS for their research question. Different RRS setups (i.e. various species and libraries constructed via different protocols, enzymes and size selection windows) were successfully sequenced together on one HiSeq lane. Most individuals included in this multi-library-multiplex received adequate sequencing effort, which has been problematic in other studies that pooled individuals directly after ligation (83). From our experience (including this and previous studies in our laboratory; see e.g. 56–58) it seems that careful, repeated quantification and standardization of DNA from every individual before and after PCR are key to achieve equivalent sequencing effort across individuals. A pilot sequencing experiment can then yield valuable insights before proceeding with sequencing at a larger scale. Here, more loci than expected were assembled in most taxa (ostracods, bivalves, sea stars) at sufficiently high per locus coverage. This highlights the value of choosing parameters conservatively, e.g. under-rather than overestimating the number of sequencing reads. The fish library yielded fewer loci than expected at higher coverage. Pooling more individuals, increasing the size window, or changing the restriction enzyme setup altogether including new optimization are future options to further optimize this project, although the current setup also yields useful data. The bird library produced coverage that is directly at the advised limit of 10× (42). This may be partly related to low quality input DNA, which was mostly extracted from feathers. Alternative sampling and/or DNA extraction protocols and further testing are needed before sequencing full scale libraries for snow petrels. Overall, a few key properties determine the feasibility and cost of RRS in non-model organisms.

### 4.1 Predictability of reduced representation experiments

Planning a genome reduction through restriction enzyme digestion starts with an imperative question: how large is the target genome? Non-model species often lack information on genome size, which complicates RRS optimization (97). If genome size appears relatively conserved across species within a taxonomic class (e.g. Asteroidea), it can be assumed that the species of interest from this class has similar genome size. Some imprecisions regarding the exact size have only limited effects on overall accuracy. Alternatively, in other groups, such as amphipods, genome size is highly variable, spanning two orders of magnitude (55,94,98). In this case, using an inaccurate genome size estimate has the potential to dramatically impact the parameters one aims to optimize. In addition, very large genomes are often highly repetitive, which significantly hampers downstream bioinformatics and population genomics (69,99,100). Therefore, with the current state of knowledge, we opted to exclude amphipods from our trial RRS libraries. Estimating genome size with flow cytometry or conducting a series of test libraries could be alternative ways forward.

For ostracods, bivalves, sea stars, and birds more loci were found than expected. This might indicate that genome sizes were consistently larger than expected. Another, likely explanation is that the enzymes used (*ApeKI, PstI*) produce more fragments than *in silico* digestions predicted. For example, the number of fragments resulting from four base cutters may be more difficult to predict as they sometimes produce so many fragments that effectively the entire genome would be sequenced (97). The five-base recognition site of *ApeKI* features a degenerate base, which may have a similar effect. The methylation sensitivity of *ApeKI* may also provide more genomic markers in genic regions (101). It is unclear, however, how general this prediction holds across metazoans. Finally, some of the excess loci recovered may be artefacts from library preparation, PCR duplicates, or incorrect locus assembly (15). Rigorous downstream filtering and/or comparison of several, differently filtered datasets may help determine the true biological signal. Whatever the reason, the higher-than-expected number of loci still lead to sufficient coverage, except in the bird library. The latter is likely related to low quality/quantity of input DNA. Few bird samples were available, some only as feathers, which yielded very little DNA. Whole genome amplification (WGA) could be an option to increase yield for RRS as successfully applied in ostracods (this study) and insects (102).

Finally, even with reliable genome size estimates and well-tested enzymes, the empirical results may differ from *in silico* expectations. In *Trematomus* fishes, approximately half of the expected sites were found, despite well-known genome size (66–68). Genomic architecture may play an important role in affecting the number of cut sites per restriction enzyme. We used the draft genome of a related species from the same family to estimate the number of fragments. The endemic Antarctic notothenioid fishes, however, are characterized by frequent chromosomal rearrangements and large numbers of transposable elements (66,103,104). The genus *Trematomus* constitutes an example of a relatively recent marine adaptive radiation (105,106). Therefore, in this particular case, the genome of a closely related species may provide relatively poor accuracy for cut site estimations.

We have tested various enzymes and enzyme combinations that have been successfully used in RRS studies (Table 1). Yet, many previous studies achieved overall relatively little marker density, which is problematic if looking for genome-wide adaptation patterns (40). With increasing output of sequencers, aiming at higher marker density is not an unachievable goal. Genome size, restriction enzyme characteristics and genomic complexity influence the predictability. Altogether, our results highlight the importance of conducting test libraries before embarking on larger, multi-library sequencing projects. In our case, *ApeKI* together with a narrow size window seems robust and powerful to create many genomic fragments (and thus sufficiently high marker density) across taxa with small to medium genome size. Using the same restriction enzyme for several projects drastically reduces cost as the same custom-made barcodes and adaptors can be used.

### 4.2 Decision making for population genomics

As we illustrated here, there are many experimental choices that may lead to inefficient or “broken” (40) RRS experiments. Given the publication bias towards successful applications (107), it is likely that a large number of unsuccessful applications of this technology to non-model species exist. It is crucially important that researchers actively engage in the decision-making process when choosing restriction enzymes, size selection windows, and the number of individuals to be pooled per sequencing lane. Furthermore, the research objectives and budget should be critically evaluated and matched. In other words, investigating genome-wide polygenic adaptation patterns in a non-model species with large, complex genomes may simply not be feasible on a small budget. The number of individuals to be included is another aspect that weighs in on these considerations. In situations where sampling is not restricted, inferences of spatial genetic structure for example may benefit more from wider geographic sampling coverage than from higher marker density. If sampling more localities is unfeasible as may be the case in the Antarctic realm, it can be beneficial to instead invest in high density sequencing (as in several markers per linkage group). With sufficient genome coverage even advanced coalescent modeling is possible using RRS data (108).

We recommend following a few guiding principles when planning RRS for population genomics (but see also e.g. 5,42,88). First, clear targets with respect to the number of individuals to be screened in a project (and/or in follow-up projects) and the marker density needed for the research objective should be defined. Second, *in silico* estimations of how these targets can be reached and approximations of the associated costs should be obtained. The number of markers and individuals must be matched to reach a certain coverage (e.g. an average target of 30×). Subsequently, it is useful to briefly evaluate the trade-offs and benefits of RRS and other methods. If a promising combination of RRS method, enzyme, size selection, and sequencing effort is found, it is often worthwhile to conduct a pilot experiment before running the full sequencing experiment. However, it is also advisable to stick to one approach afterwards and not change for example the sequencing platform, the size window or other properties of the setup that will otherwise reduce comparability between datasets.

### 4.3 Alternative approaches

In some cases, RRS might not be the right choice for molecular ecological research. A plethora of other genomic or genetic methods exists, which may offer more appropriate cost-benefit ratios. SNP genotyping arrays are a common and highly reproducible alternative, but usually only for species with more genomic resources (which exist also for some Antarctic taxa; see 76). Similarly, whole genome resequencing is providing the most extensive datasets which can be used for a wide range of analyses (11,12,110). However, this is still too costly for many research projects, especially if information across many individuals and/or localities is needed. Another option is to focus on the expressed part of the genome and use a form of sequence capture enrichment (e.g. 64,78,79) or RNAseq (113), or both (114,115). These approaches are versatile and can provide valuable information, even for museum samples (116,117). However, substantial expertise and prior investment the development of custom methods is necessary for species that have not been investigated yet. With a limited budget and research objectives that do not depend on whole genome scans for selection, more classical molecular approaches are sometimes a good alternative. Nuclear microsatellite markers remain powerful to describe population structure and can be multiplexed and screened in large numbers. These markers can also benefit from high-throughput sequencing (118,119). Mitogenome sequencing and assembly using long-range PCR is another useful approach, particularly for phylogeographic applications (120,121). The amphipod and bird species evaluated here may currently be more amenable to such methods instead of RRS.

### 4.4 Conclusions

An extensive evaluation and optimization protocol allowed us to identify whether RRS is a suitable option for population genomics in a range of Antarctic animals. We have achieved promising results in some classes (ostracods, bivalves, sea stars, and fishes) that will be further developed soon. In other cases (amphipods and birds/degraded samples) alternative strategies such as mitogenome, capture sequencing or microsatellites seem more appropriate. The detailed considerations outlined here are a guideline for researchers to make informed decisions about the use of RRS or alternative methods. This is particularly important for species where genomic information remains scarce.

## Declarations

### • Ethics approval and consent to participate

Animal tissues were sampled following internationally recognized CCAMLR CEMP standard methods and were permitted under the Antarctic Marine Living Resources Act.

### • Consent for publication

Not applicable.

### • Availability of data and materials

The datasets supporting the conclusions of this article are available in the NCBI’s Sequence Read Archive (SRA) repository, BioProject ID PRJNA674352, reviewer link: https://dataview.ncbi.nlm.nih.gov/object/PRJNA674352?reviewer=dmj6c5816761lpqn2d1oe69qhs, and in the Zenodo repository, https://doi.org/10.5281/zenodo.3267164.

### • Authors’ contributions

HC and IS conceived the study. BH, FH, HC, QJ, FP, HR, MV and IS conducted laboratory work. HC analyzed the data and wrote the first draft of the manuscript. All authors contributed samples, intellectual input during project meetings and comments on the manuscript.

## • Acknowledgements

We are very grateful to everyone that contributed to field work and provided samples. In particular, we thank S. Brandão for ostracod DNA and H. Griffiths and K. Linse (BAS), C. Held (AWI), S. Mills and D. Macpherson (NIWA), H. Baird and G. Johnstone (AAD), and the MNHN in Paris for amphipod samples. We thank M. Clark (BAS) and the scientific staff involved for the generous collection of bivalve samples from Rothera station. We thank the captains, crew and science leaders who contributed to the collection of sea stars from RRS *James Clark Ross* (JR15005) and RV *Polarstern* (PS77, PS81). We thank the scientific team of IPEV (Institut Polaire Français Paul-Emile Victor) program 1124 REVOLTA (Radiation EVOLutives en Terre Adélie; chief scientists M. Eléaume, N. Améziane and G. Lecointre). For fishes, we thank K. Roche and P. Jurajda who provided fin clips from James Ross Island. The toothfish fishery observers and S. Parker (NIWA) are acknowledged for fish samples from the Ross Sea. We thank G.E. Maes and K. Herten from the KU Leuven Genomics Core for technical support and advice.

## • Authors’ information

Not applicable.

## List of Figures

**Figure 3.**
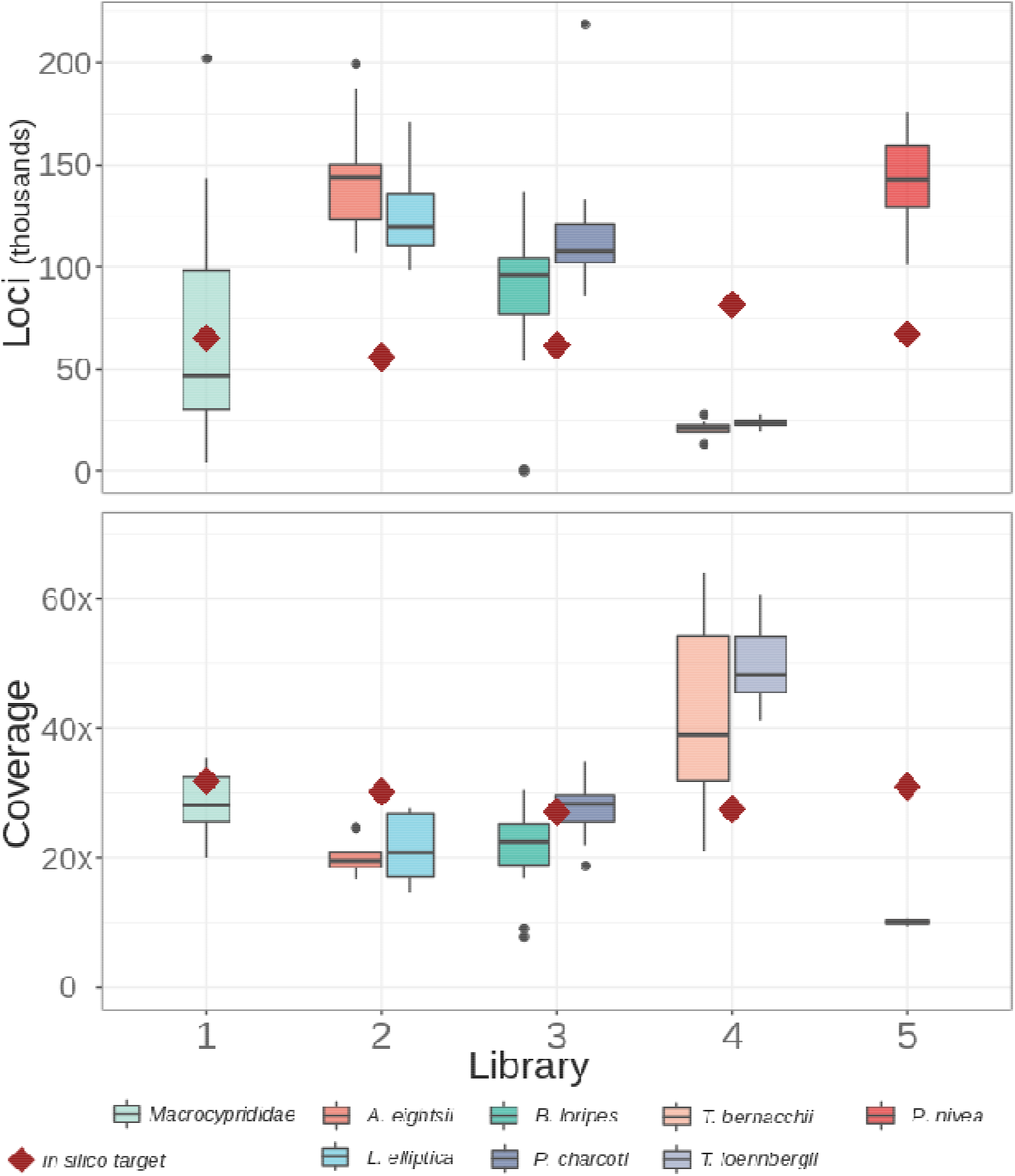
The number of loci and coverage as estimated and realized in five test libraries. *In silico* estimates (dark red diamonds) and empirical values from reduced representation sequencing (RRS) libraries containing DNA from eight target species are shown. Boxplots show the median, quartiles, and outliers across individuals (N=8-14). Libraries were prepared as listed in Table 4.

## List of Supplemental Information

**Additional File 1. DOCX. Samples used for reduced representation sequencing (RRS) optimization.** DNA from these samples was used for empirical restriction enzyme digestions with different enzymes (single digest *EcoRI, PstI, MspI*, or double digest *EcoRI-MspI*) and for RRS pilot libraries. Some samples were extracted twice as replicates (marked as _rep in sample ID). Three samples per species (family in the case of ostracods) were used for empirical digestions. The amphipod (*C. obesa* and *E. pontomedon*) samples and one *T. loennbergii* were used for empirical digestions, but not included in any RRS library.

**Additional File 2. DOCX. *In silico* estimates of the number of fragments.** Estimates were produced through *in silico* restriction enzyme digestions for reduced representation sequencing (RRS) optimized for approximately 30× coverage. The number of fragments depends on the restriction enzyme/combination, the size window, the assumed genome size, and the reference genome used for *in silico* digestion. Reference genomes of related species were used as well as simulated genomes; in this case the size and GC content used to simulate the genomes are listed. The number of fragments were extrapolated to the assumed genome size. Only two different enzyme and size selection setups per target species are listed here (for RRS setups optimized for HiSeq 2500 or HiSeq 4000 sequencing runs, respectively; the same as in Table 3, Table 4, Additional File 4); further estimates can be found in spreadsheets available at https://doi.org/10.5281/zenodo.3267164.

**Additional File 3. DOCX. Comparisons of empirical and *in silico* restriction enzyme digestions.** Empirical Bioanalyzer results (left figure panels) with digested DNA are shown as concentration over fragment size and estimated loci numbers over locus size from *in silico* digestions (right figure panels) for all target taxa except fish (these are shown in Fig. 2).

**Additional File 4. DOCX. Reduced representation sequencing (RRS) setups for seven individually optimized protocols.** These setups were optimized in order to be run on a HiSeq 4000 platform (Illumina). The choice of restriction enzyme(s) and size window was optimized to obtain approximately 30× coverage (or half that in a worst-case scenario) with the assumed genome size (conservatively estimated based on available information, see Table 1). Marker density was estimated as a comparable measure to the metastudy by Lowry et al. (40).

**Additional File 5. DOCX. Reduced representation sequencing (RRS) laboratory protocol based on the protocol from Peterson et al. (17).** The protocol is scaled for use with 192 samples and with restriction enzymes *EcoRI* and *MspI*; the reagent volumes can be scaled down/up to suit other sample numbers; if other enzymes are used, the respective reaction conditions must be adjusted.

**Additional File 6. DOCX. Reduced representation sequencing (RRS) laboratory protocol based on the protocol from Elshire et al. (20).** The protocol is scaled for use with 192 samples and with restriction enzymes *PstI* or *ApeKI*; the reagent volumes can be scaled down/up to suit other sample numbers; if other enzymes are used, the respective reaction conditions must be adjusted.

**Additional File 7. DOCX. Results from parameter optimization for *de novo* assembly and genotyping.** Eight parameter optimization series were conducted following Rochette & Catchen (42) to identify optimal parameters to genotype reduced representation sequencing (RRS) data with Stacks v2.4 (15); one test series for each species/species complex. The Stacks parameter m was kept constant (m = 3), while parameters M and n were varied together from 1 to 9. Subsequently, only loci present in 80 % of the samples were retained and for each M=n parameter the number of loci and polymorphic loci was plotted, as well as the proportion of these loci containing 0 to 10 or >10 SNPs. In ostracods, the library contained DNA from a species-complex, resulting in very few shared loci across 80 % of the samples. Therefore, in this case results based on loci shared by 50 % of samples are shown. Optimal M=n values were decided in all cases with this information (and reported in Table 4). Note, however, that it is impossible to make absolute calls regarding the ideal value.

