## Supplemental Information for "Facilitating population genomics of non-model organisms through optimized experimental design for reduced representation sequencing"

*Correspondence: Henrik Christiansen

**Additional File 1. Samples used for reduced representation sequencing (RRS) optimization.** DNA from these samples was used for empirical restriction enzyme digestions with different enzymes (single digest *EcoRI*, *PstI*, *MspI*, or double digest *EcoRI-MspI*) and for RRS pilot libraries. Some samples were extracted twice as replicates (marked as _rep in sample ID). Three samples per species (family in the case of ostracods) were used for empirical digestions. The amphipod (*C. obesa* and *E. pontomedon*) samples and one *T. loennbergii* were used for empirical digestions, but not included in any RRS library.

| **Species** | **Sample ID** | **Origin** | **Empirical Digestion** | **RRS Library** |
| --- | --- | --- | --- | --- |
| *Macropyxis hornei* | 280 | ANT XIX-3, St. 46-7-S | *EcoRI*, *MspI* | 1 |
| *Macrocyprina rocas* | 340 | Buzios | *EcoRI*, *MspI* | 1 |
| *Macroscapha falcis* | 176 | ANT XXII-3, St. 74-6-S | *EcoRI*, *MspI* | 1 |
| *Macroscapha falcis* | 187 | ANT XXII-3, St. 74-6-E | *-* | 1 |
| *Macroscapha solecavai* | 223 | ANT XXII-3, St. 151-7-E | *-* | 1 |
| *Macroscapha falcis* | 186 | ANT XXII-3, St. 74-6-E | *-* | 1 |
| *Macroscapha opaca* | 240 | ANT XXII-3, St. 153-7-S | *-* | 1 |
| *Macroscapha solecavai* | 226 | ANT XXII-3, St. 151-7-E | *-* | 1 |
| *Macroscapha opaca* | 240_rep | ANT XXII-3, St. 153-7-S | *-* | 1 |
| *Macroscapa solecavai* | 226_rep | ANT XXII-3, St. 151-7-E | *-* | 1 |
| *Charcotia obesa* | A77 | ANTXXIX-3 PS81,  St. 162-7 | *EcoRI*, *MspI* | - |
| *Charcotia obesa* | A78 | ANTXXIX-3 PS81,  St. 162-7 | *EcoRI*, *MspI* | - |
| *Charcotia obesa* | A79 | ANTXXIX-3 PS81,  St. 162-7 | *EcoRI*, *MspI* | - |
| *Eusirus pontomedon* | HE10 | ANTXXIX-3 PS81,  St. 227-2 | *EcoRI*, *MspI* | - |
| *Eusirus pontomedon* | HE13 | ANTXXIX-3 PS81,  St. 227-2 | *EcoRI*, *MspI* | - |
| *Eusirus pontomedon* | HE14 | ANTXXIX-3 PS81,St. 227-2 | *EcoRI*, *MspI* | - |
| *Laternula elliptica* | 4C | - | *EcoRI*, *PstI*, *MspI* | 2 |
| *Laternula elliptica* | 5C | - | *EcoRI*, *PstI*, *MspI* | 2 |
| *Laternula elliptica* | 6C | - | *EcoRI*, *PstI*, *MspI* | 2 |
| *Laternula elliptica* | KGI18 | - | *-* | 2 |
| *Laternula elliptica* | KGI11 | - | *-* | 2 |
| *Laternula elliptica* | R5 | - | *-* | 2 |
| *Laternula elliptica* | R6 | - | *-* | 2 |
| *Laternula elliptica* | R7 | - | *-* | 2 |
| *Laternula elliptica* | R6_rep | - | *-* | 2 |
| *Laternula elliptica* | R7_rep | - | *-* | 2 |
| *Aequiyoldia eightsii* | 1C | - | *EcoRI*, *PstI*, *MspI* | 2 |
| *Aequiyoldia eightsii* | 2C | - | *EcoRI*, *PstI*, *MspI* | 2 |
| *Aequiyoldia eightsii* | 3C | - | *EcoRI*, *PstI*, *MspI* | 2 |
| *Aequiyoldia eightsii* | R10 | - | *-* | 2 |
| *Aequiyoldia eightsii* | R11 | - | *-* | 2 |
| *Aequiyoldia eightsii* | R12 | - | *-* | 2 |
| *Aequiyoldia eightsii* | R27 | - | *-* | 2 |
| *Aequiyoldia eightsii* | R28 | - | *-* | 2 |
| *Aequiyoldia eightsii* | KGI2 | - | *-* | 2 |
| *Aequiyoldia eightsii* | KGI5 | - | *-* | 2 |
| *Aequiyoldia eightsii* | R27_rep | - | *-* | 2 |
| *Aequiyoldia eightsii* | R28_rep | - | *-* | 2 |
| *Bathybiaster loripes* | Bat004 | Proteker II | *-* | 3 |
| *Bathybiaster loripes* | Bat062 | CEAMARC | *-* | 3 |
| *Bathybiaster loripes* | Bat076 | LASSO_ANTXXIX/3 | *EcoRI*, *PstI*, *MspI* | 3 |
| *Bathybiaster loripes* | Bat095 | ANT XXVII/3 (CAMBIO) | *-* | 3 |
| *Bathybiaster loripes* | Bat096 | ANT XXVII/3 (CAMBIO) | *EcoRI*, *PstI*, *MspI* | 3 |
| *Bathybiaster loripes* | Bat152 | ANT XXVII/3 (CAMBIO) | *EcoRI*, *PstI*, *MspI* | 3 |
| *Bathybiaster loripes* | Bat156 | JR230 | *-* | 3 |
| *Bathybiaster loripes* | Bat157 | JR275 | *-* | 3 |
| *Bathybiaster loripes* | Bat164 | POKER II | *-* | 3 |
| *Bathybiaster loripes* | Bat184 | POKER II | *-* | 3 |
| *Bathybiaster loripes* | Bat004_rep | Proteker II | *-* | 3 |
| *Bathybiaster loripes* | Bat062_rep | CEAMARC | *-* | 3 |
| *Psilaster charcoti* | Psi002 | JR15005 | *EcoRI*, *PstI*, *MspI* | 3 |
| *Psilaster charcoti* | Psi003 | JR15005 | *EcoRI*, *PstI*, *MspI* | 3 |
| *Psilaster charcoti* | Psi008 | JR15005 | *-* | 3 |
| *Psilaster charcoti* | Psi036 | REVOLTA1 | *EcoRI*, *PstI*, *MspI* | 3 |
| *Psilaster charcoti* | Psi037 | REVOLTA1 | *-* | 3 |
| *Psilaster charcoti* | Psi039 | JR275 | *-* | 3 |
| *Psilaster charcoti* | Psi040 | JR275 | *-* | 3 |
| *Psilaster charcoti* | Psi048 | CEAMARC | *-* | 3 |
| *Psilaster charcoti* | Psi063 | CEAMARC | *-* | 3 |
| *Psilaster charcoti* | Psi075 | CEAMARC | *-* | 3 |
| *Psilaster charcoti* | Psi153 | JR275 | *-* | 3 |
| *Psilaster charcoti* | Psi155 | JR275 | *-* | 3 |
| *Psilaster charcoti* | Psi164 | JR15005 | *-* | 3 |
| *Psilaster charcoti* | Psi215 | JR308 | *-* | 3 |
| *Psilaster charcoti* | Psi037_rep | REVOLTA1 | *-* | 3 |
| *Psilaster charcoti* | Psi039_rep | JR275 | *-* | 3 |
| *Trematomus bernacchii* | JRI_02 | *see* Jurajda et al. (126) | *-* | 4 |
| *Trematomus bernacchii* | JRI_03 | *see* Jurajda et al. (126) | *-* | 4 |
| *Trematomus bernacchii* | JRI_04 | *see* Jurajda et al. (126) | *-* | 4 |
| *Trematomus bernacchii* | JRI_05 | *see* Jurajda et al. (126) | *-* | 4 |
| *Trematomus bernacchii* | JRI_06 | *see* Jurajda et al. (126) | *EcoRI*, *ApeKI*, *EcoRI-MspI* | 4 |
| *Trematomus bernacchii* | JRI_07 | *see* Jurajda et al. (126) | *-* | 4 |
| *Trematomus bernacchii* | JRI_08 | *see* Jurajda et al. (126) | *EcoRI*, *ApeKI*, *EcoRI-MspI* | 4 |
| *Trematomus bernacchii* | JRI_09 | *see* Jurajda et al. (126) | *-* | 4 |
| *Trematomus bernacchii* | JRI_10 | *see* Jurajda et al. (126) | *-* | 4 |
| *Trematomus bernacchii* | JRI_11 | *see* Jurajda et al. (126) | *EcoRI*, *ApeKI*, *EcoRI-MspI* | 4 |
| *Trematomus bernacchii* | JRI_03_rep | *see* Jurajda et al. (126) | *-* | 4 |
| *Trematomus bernacchii* | JRI_04_rep | *see* Jurajda et al. (126) | *-* | 4 |
| *Trematomus loennbergii* | ROS_1352 | RSSS 2016 | *-* | 4 |
| *Trematomus loennbergii* | ROS_1353 | RSSS 2016 | *-* | 4 |
| *Trematomus loennbergii* | ROS_1354 | RSSS 2016 | *-* | 4 |
| *Trematomus loennbergii* | ROS_1417 | RSSS 2016 | *-* | 4 |
| *Trematomus loennbergii* | ROS_1418 | RSSS 2016 | *EcoRI*, *ApeKI*, *EcoRI-MspI* | 4 |
| *Trematomus loennbergii* | ROS_1419 | RSSS 2016 | *-* | 4 |
| *Trematomus loennbergii* | ROS_1420 | RSSS 2016 | *-* | 4 |
| *Trematomus loennbergii* | ROS_1421 | RSSS 2016 | *-* | 4 |
| *Trematomus loennbergii* | ROS_1484 | RSSS 2016 | *-* | 4 |
| *Trematomus loennbergii* | ROS_1485 | RSSS 2016 | *EcoRI*, *ApeKI*, *EcoRI-MspI* | 4 |
| *Trematomus loennbergii* | ROS_1487 | RSSS 2016 | *EcoRI*, *ApeKI*, *EcoRI-MspI* | - |
| *Trematomus loennbergii* | ROS_1352_rep | RSSS 2016 | *-* | 4 |
| *Trematomus loennbergii* | ROS_1353_rep | RSSS 2016 | *-* | 4 |
| *Pagodroma nivea* | 1 | BAS, Rothera Point | *-* | 5 |
| *Pagodroma nivea* | 2 | BAS, Storm Ridge | *EcoRI*, *PstI*, *MspI* | 5 |
| *Pagodroma nivea* | 3 | BAS, Storm Ridge | *EcoRI*, *PstI*, *MspI* | 5 |
| *Pagodroma nivea* | 4 | BAS, Signy Island | *EcoRI*, *PstI*, *MspI* | 5 |
| *Pagodroma nivea* | 5 | BAS, Signy Island | *EcoRI*, *PstI*, *MspI* | 5 |
| *Pagodroma nivea* | BEL-G05 | Ut 005 | *-* | 5 |
| *Pagodroma nivea* | BEL-G81 | Ta 081 | *-* | 5 |
| *Pagodroma nivea* | BEL-G20 | Pi 020 | *-* | 5 |
| *Pagodroma nivea* | BEL-G05_rep1 | Ut 005 | *-* | 5 |
| *Pagodroma nivea* | BEL-G05_rep2 | Ut 005 | *-* | 5 |

**Additional File 2. *In silico* estimates of the number of fragments.** Estimates were produced through *in silico* restriction enzyme digestions for reduced representation sequencing (RRS) optimized for approximately 30× coverage. The number of fragments depends on the restriction enzyme/combination, the size window, the assumed genome size, and the reference genome used for *in silico* digestion. Reference genomes of related species were used as well as simulated genomes; in this case the size and GC content used to simulate the genomes are listed. The number of fragments were extrapolated to the assumed genome size. Only two different enzyme and size selection setups per target species are listed here (for RRS setups optimized for HiSeq 2500 or HiSeq 4000 sequencing runs, respectively; the same as in Table 3, Table 4, Additional File 4); further estimates can be found in spreadsheets available at <https://doi.org/10.5281/zenodo.3267164>.

| Class | Target Species | Restriction Enzyme (Combination) | Size Window (bp) | Assumed Genome Size (Mb) | Reference genome (GC content) | Number of fragments |
| --- | --- | --- | --- | --- | --- | --- |
| *Ostracoda* | Macrocyprididae | *ApeKI* | 200-350 | 250 | *C. torosa* | 65,244 |
| *Ostracoda* | Macrocyprididae | *ApeKI* | 200-350 | 250 | *D. pulex* | 53,331 |
| *Ostracoda* | Macrocyprididae | *ApeKI* | 200-350 | 250 | *A. tonsa* | 15,122 |
| *Ostracoda* | Macrocyprididae | *ApeKI* | 200-350 | 250 | *P. hawaiensis* | 14,927 |
| *Ostracoda* | Macrocyprididae | *ApeKI* | 200-350 | 250 | 100 Mb (43.9) | 42,545 |
| *Ostracoda* | Macrocyprididae | *ApeKI* | 200-350 | 250 | 500 Mb (43.9) | 42,425 |
| *Ostracoda* | Macrocyprididae | *ApeKI* | 250-500 | 250 | *C. torosa* | 88,550 |
| *Ostracoda* | Macrocyprididae | *ApeKI* | 250-500 | 250 | *D. pulex* | 70,618 |
| *Ostracoda* | Macrocyprididae | *ApeKI* | 250-500 | 250 | *A. tonsa* | 22,105 |
| *Ostracoda* | Macrocyprididae | *ApeKI* | 250-500 | 250 | *P. hawaiensis* | 19,078 |
| *Ostracoda* | Macrocyprididae | *ApeKI* | 250-500 | 250 | 100 Mb (43.9) | 62,688 |
| *Ostracoda* | Macrocyprididae | *ApeKI* | 250-500 | 250 | 500 Mb (43.9) | 62,476 |
| *Malacostraca* | *Charcotia obesa* | *SbfI_MspI* | 200-330 | 27,000 | *H. azteca* | 64,094 |
| *Malacostraca* | *Charcotia obesa* | *SbfI_MspI* | 200-330 | 27,000 | *P. hawaiensis* | 25,118 |
| *Malacostraca* | *Charcotia obesa* | *SbfI_MspI* | 200-330 | 27,000 | *E. perdentatus*^†^ | 10,984 |
| *Malacostraca* | *Charcotia obesa* | *SbfI_MspI* | 200-330 | 27,000 | 10,000 Mb (38.5) | 31,590 |
| *Malacostraca* | *Charcotia obesa* | *SbfI_MspI* | 200-330 | 27,000 | 30,000 Mb (40.8) | 44,820 |
| *Malacostraca* | *Charcotia obesa* | *SbfI_MspI* | 250-450 | 27,000 | *H. azteca* | 91,927 |
| *Malacostraca* | *Charcotia obesa* | *SbfI_MspI* | 250-450 | 27,000 | *P. hawaiensis* | 41,576 |
| *Malacostraca* | *Charcotia obesa* | *SbfI_MspI* | 250-450 | 27,000 | *E. perdentatus*^†^ | 13,863 |
| *Malacostraca* | *Charcotia obesa* | *SbfI_MspI* | 250-450 | 27,000 | 10,000 Mb (38.5) | 50,760 |
| *Malacostraca* | *Charcotia obesa* | *SbfI_MspI* | 250-450 | 27,000 | 30,000 Mb (40.8) | 61,155 |
| *Malacostraca* | *Eusirus pontomedon* | *EcoRI_SphI* | 200-260 | 7,000 | *H. azteca* | 63,572 |
| *Malacostraca* | *Eusirus pontomedon* | *EcoRI_SphI* | 200-260 | 7,000 | *P. hawaiensis* | 12,800 |
| *Malacostraca* | *Eusirus pontomedon* | *EcoRI_SphI* | 200-260 | 7,000 | *E. pontomedon*^†^ | 10,986 |
| *Malacostraca* | *Eusirus pontomedon* | *EcoRI_SphI* | 200-260 | 7,000 | 10,000 Mb (38.5) | 41,580 |
| *Malacostraca* | *Eusirus pontomedon* | *EcoRI_SphI* | 200-260 | 7,000 | 30,000 Mb (40.8) | 45,325 |
| *Malacostraca* | *Eusirus pontomedon* | *EcoRI_SphI* | 250-350 | 7,000 | *H. azteca* | 101,900 |
| *Malacostraca* | *Eusirus pontomedon* | *EcoRI_SphI* | 250-350 | 7,000 | *P. hawaiensis* | 22,047 |
| *Malacostraca* | *Eusirus pontomedon* | *EcoRI_SphI* | 250-350 | 7,000 | *E. pontomedon*^†^ | 18,000 |
| *Malacostraca* | *Eusirus pontomedon* | *EcoRI_SphI* | 250-350 | 7,000 | 10,000 Mb (38.5) | 63,070 |
| *Malacostraca* | *Eusirus pontomedon* | *EcoRI_SphI* | 250-350 | 7,000 | 30,000 Mb (40.8) | 71,225 |
| *Bivalvia* | *Laternula elliptica* & *Aequiyoldia eightsii* | *ApeKI* | 200-260 | 3,000 | *C. gigas* | 69,027 |
| *Bivalvia* | *Laternula elliptica* & *Aequiyoldia eightsii* | *ApeKI* | 200-260 | 3,000 | *P. imbricata* | 64,486 |
| *Bivalvia* | *Laternula elliptica* & *Aequiyoldia eightsii* | *ApeKI* | 200-260 | 3,000 | *B. platifrons* | 53,399 |
| *Bivalvia* | *Laternula elliptica* & *Aequiyoldia eightsii* | *ApeKI* | 200-260 | 3,000 | 1,000 Mb (35.3) | 59,400 |
| *Bivalvia* | *Laternula elliptica* & *Aequiyoldia eightsii* | *ApeKI* | 200-260 | 3,000 | 5,000 Mb (34.2) | 45,480 |
| *Bivalvia* | *Laternula elliptica* & *Aequiyoldia eightsii* | *ApeKI* | 250-350 | 3,000 | *C. gigas* | 105,349 |
| *Bivalvia* | *Laternula elliptica* & *Aequiyoldia eightsii* | *ApeKI* | 250-350 | 3,000 | *P. imbricata* | 102,333 |
| *Bivalvia* | *Laternula elliptica* & *Aequiyoldia eightsii* | *ApeKI* | 250-350 | 3,000 | *B. platifrons* | 83,580 |
| *Bivalvia* | *Laternula elliptica* & *Aequiyoldia eightsii* | *ApeKI* | 250-350 | 3,000 | 1,000 Mb (35.3) | 96,600 |
| *Bivalvia* | *Laternula elliptica* & *Aequiyoldia eightsii* | *ApeKI* | 250-350 | 3,000 | 5,000 Mb (34.2) | 75,420 |
| *Asteroidea* | *Bathybiaster loripes & Psilaster charcoti* | *ApeKI* | 200-300 | 500 | *A. planci* | 76,988 |
| *Asteroidea* | *Bathybiaster loripes & Psilaster charcoti* | *ApeKI* | 200-300 | 500 | *P. miniata* | 64,466 |
| *Asteroidea* | *Bathybiaster loripes & Psilaster charcoti* | *ApeKI* | 200-300 | 500 | *P. regularis* | 62,272 |
| *Asteroidea* | *Bathybiaster loripes & Psilaster charcoti* | *ApeKI* | 200-300 | 500 | 1,000 Mb (41.3) | 42,245 |
| *Asteroidea* | *Bathybiaster loripes & Psilaster charcoti* | *ApeKI* | 200-300 | 500 | 2,000 Mb (40.4) | 36,618 |
| *Asteroidea* | *Bathybiaster loripes & Psilaster charcoti* | *ApeKI* | 250-400 | 500 | *A. planci* | 98,911 |
| *Asteroidea* | *Bathybiaster loripes & Psilaster charcoti* | *ApeKI* | 250-400 | 500 | *P. miniata* | 79,380 |
| *Asteroidea* | *Bathybiaster loripes & Psilaster charcoti* | *ApeKI* | 250-400 | 500 | *P. regularis* | 83,222 |
| *Asteroidea* | *Bathybiaster loripes & Psilaster charcoti* | *ApeKI* | 250-400 | 500 | 1,000 Mb (41.3) | 62,144 |
| *Asteroidea* | *Bathybiaster loripes & Psilaster charcoti* | *ApeKI* | 250-400 | 500 | 2,000 Mb (40.4) | 108,693 |
| *Actinopterygii* | *Trematomus bernacchii & T. loennbergii* | *EcoRI_MspI* | 200-450 | 1,500 | *N. coriiceps* | 81,605 |
| *Actinopterygii* | *Trematomus bernacchii & T. loennbergii* | *EcoRI_MspI* | 200-450 | 1,500 | 1,000 Mb (40.8) | 213,285 |
| *Actinopterygii* | *Trematomus bernacchii & T. loennbergii* | *EcoRI_MspI* | 200-450 | 1,500 | 1,800 Mb (40.8) | 214,613 |
| *Actinopterygii* | *Trematomus bernacchii & T. loennbergii* | *EcoRI_MspI* | 200-600 | 1,500 | *N. coriiceps* | 101,138 |
| *Actinopterygii* | *Trematomus bernacchii & T. loennbergii* | *EcoRI_MspI* | 200-600 | 1,500 | 1,000 Mb (40.8) | 246,555 |
| *Actinopterygii* | *Trematomus bernacchii & T. loennbergii* | *EcoRI_MspI* | 200-600 | 1,500 | 1,800 Mb (40.8) | 247,890 |
| *Aves* | *Pagodroma nivea* | *PstI* | 200-300 | 1,500 | *F. glacialis* | 66,258 |
| *Aves* | *Pagodroma nivea* | *PstI* | 200-300 | 1,500 | 1,500 Mb (41.2) | 3,270 |
| *Aves* | *Pagodroma nivea* | *PstI* | 200-300 | 1,500 | 2,000 Mb (41.2) | 3,345 |
| *Aves* | *Pagodroma nivea* | *PstI* | 250-400 | 1,500 | *F. glacialis* | 92,422 |
| *Aves* | *Pagodroma nivea* | *PstI* | 250-400 | 1,500 | 1,500 Mb (41.2) | 5,295 |
| *Aves* | *Pagodroma nivea* | *PstI* | 250-400 | 1,500 | 2,000 Mb (41.2) | 5,040 |

^†^ no reference genome of *Eusirus pontomedon* was available, instead we here used shotgun sequencing data available from a microsatellite development project for the species

**Additional File 3. Comparisons of empirical and *in silico* restriction enzyme digestions.** Empirical Bioanalyzer results (left figure panels) with digested DNA are shown as concentration over fragment size and estimated loci numbers over locus size from *in silico* digestions (right figure panels) for all target taxa except fish (these are shown in Fig. 2).

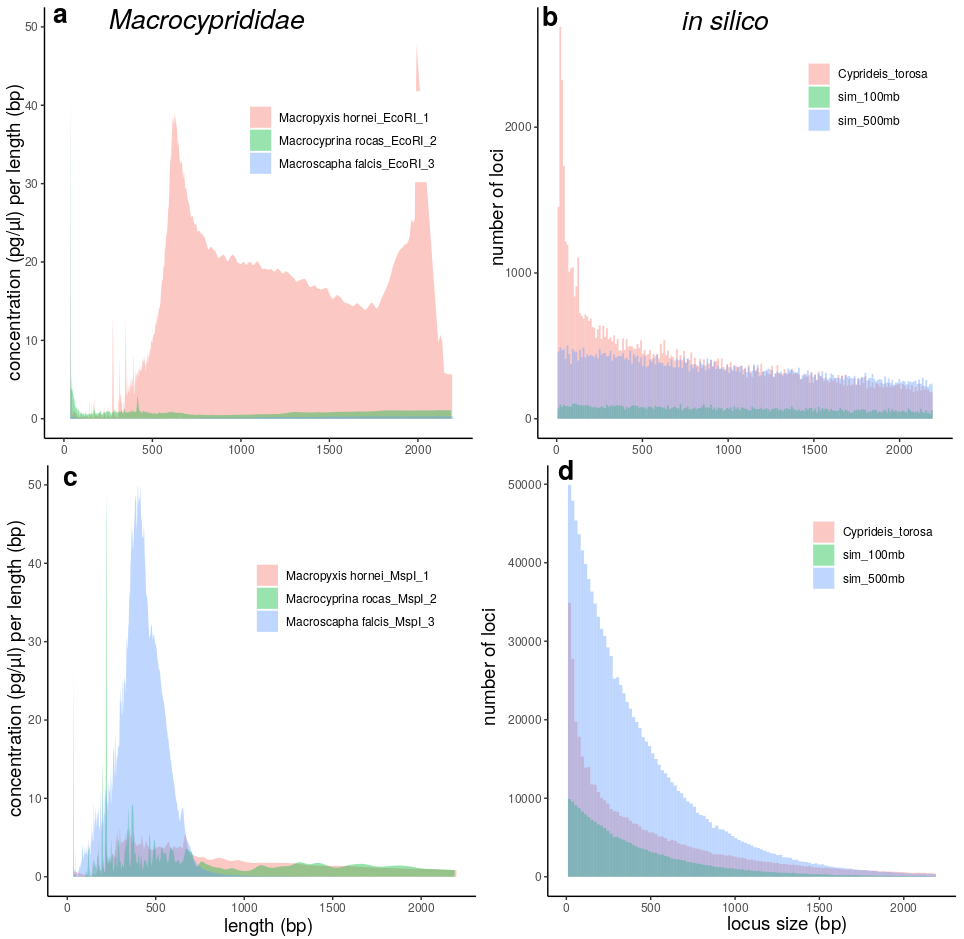

**Figure S3.1.** Empirical Bioanalyzer results with digested DNA are shown as concentration over fragment size (a, c) and estimated loci numbers over locus size from *in silico* digestions (b, d). The tests were conducted with restrictions enzymes *EcoRI* (a, b) and *MspI* (c, d). Results for the ostracod species *Macropyxis hornei*, *Macrocyprina rocas* and *Macroscapha falcis* are shown next to *in silico* estimates using a related reference genome of *Cyprideis torosa* and two simulated genomes of 100 and 500 Mb size.

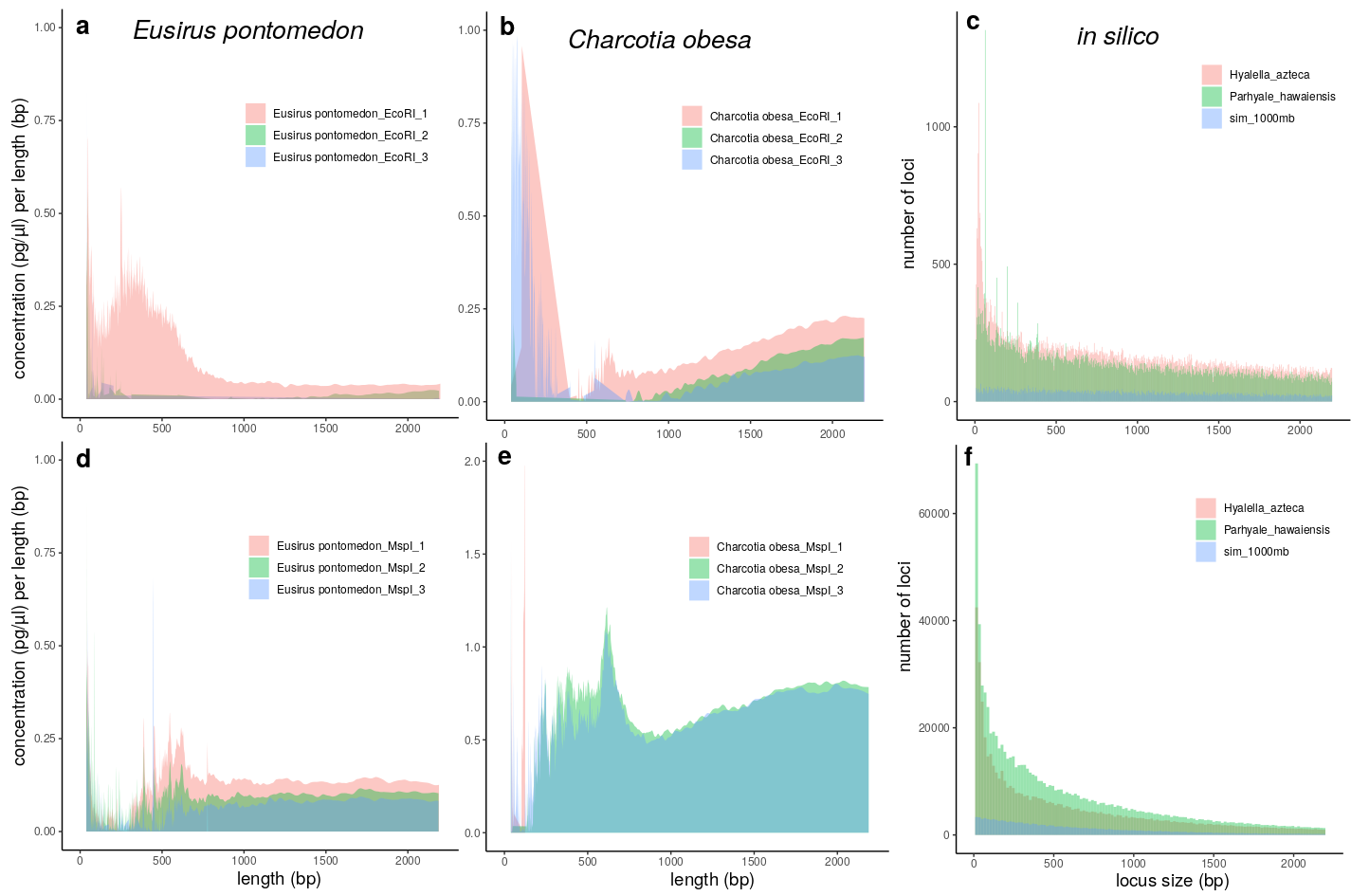

**Figure S3.2.** Empirical Bioanalyzer results with digested DNA are shown as concentration over fragment size (a, b, d, e) and estimated loci numbers over locus size from *in silico* digestions (c, f). The tests were conducted with restrictions enzymes *EcoRI* (a, b, c) and *MspI* (d, e, f). Results for the amphipods species *Eusirus pontomedon* (b, e) and *Charcotia obesa* (a, d) are shown next to *in silico* estimates using reference genomes of *Hyalella azteca*, *Parhyale hawaiensis* and one simulated genome of 1000 Mb size (note that this was the absolute size used for *in silico* computations, but resulting estimates were extrapolated to 10,000 Mb).

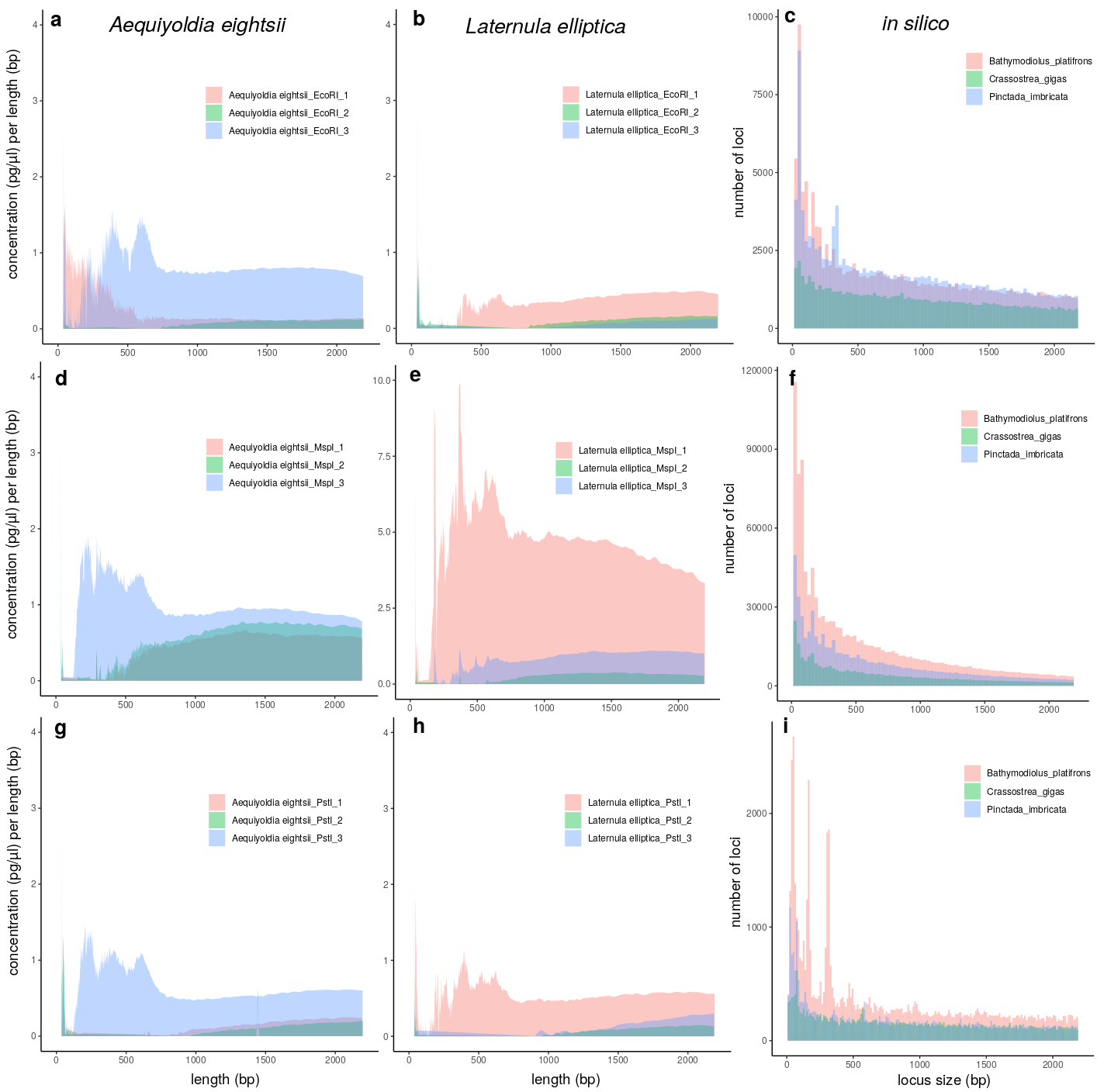

**Figure S3.3.** Empirical Bioanalyzer results with digested DNA are shown as concentration over fragment size (a, b, d, e, g, h) and estimated loci numbers over locus size from *in silico* digestions (c, f, i). The tests were conducted with restrictions enzymes *EcoRI* (a, b, c), *PstI* (d, e, f) and *MspI* (g, h, i). Results for the bivalve species *Aequiyoldia eightsii* (a, d, g) and *Laternula elliptica* (b, e, h) are shown next to *in silico* estimates using reference genomes of *Bathymodiolus platifrons, Crassostrea gigas* and *Pinctada imbricata*.

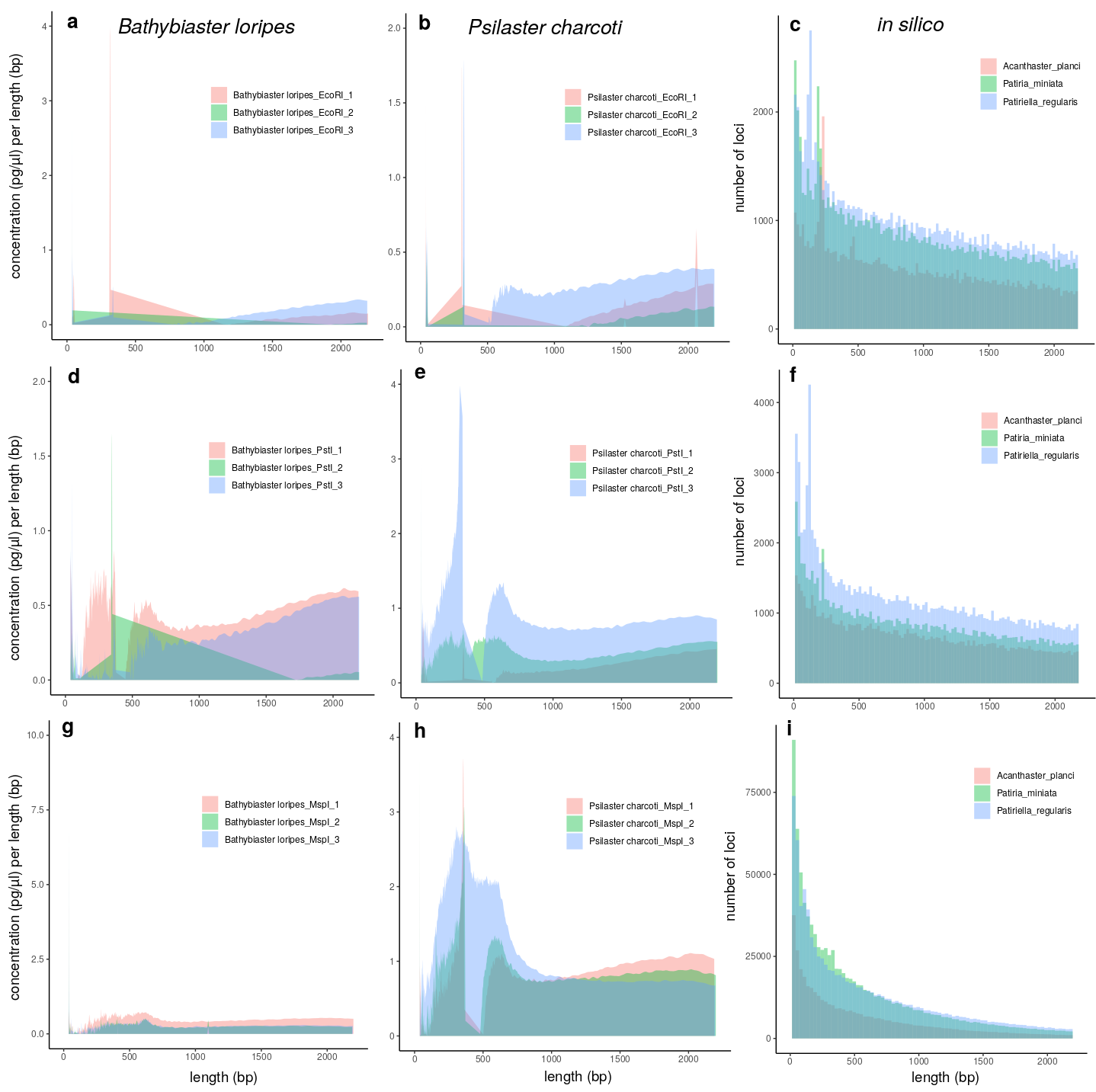

**Figure S3.4.** Empirical Bioanalyzer results with digested DNA are shown as concentration over fragment size (a, b, d, e, g, h) and estimated loci numbers over locus size from *in silico* digestions (c, f, i). The tests were conducted with restrictions enzymes *EcoRI* (a, b, c), *PstI* (d, e, f) and *MspI* (g, h, i). Results for the sea star species *Bathybiaster loripes* (a, d, g) and *Psilaster charcoti* (b, e, h) are shown next to *in silico* estimates using reference genomes of *Acanthaster planci*, *Patiria miniata*, and *Patiruella regularis*.

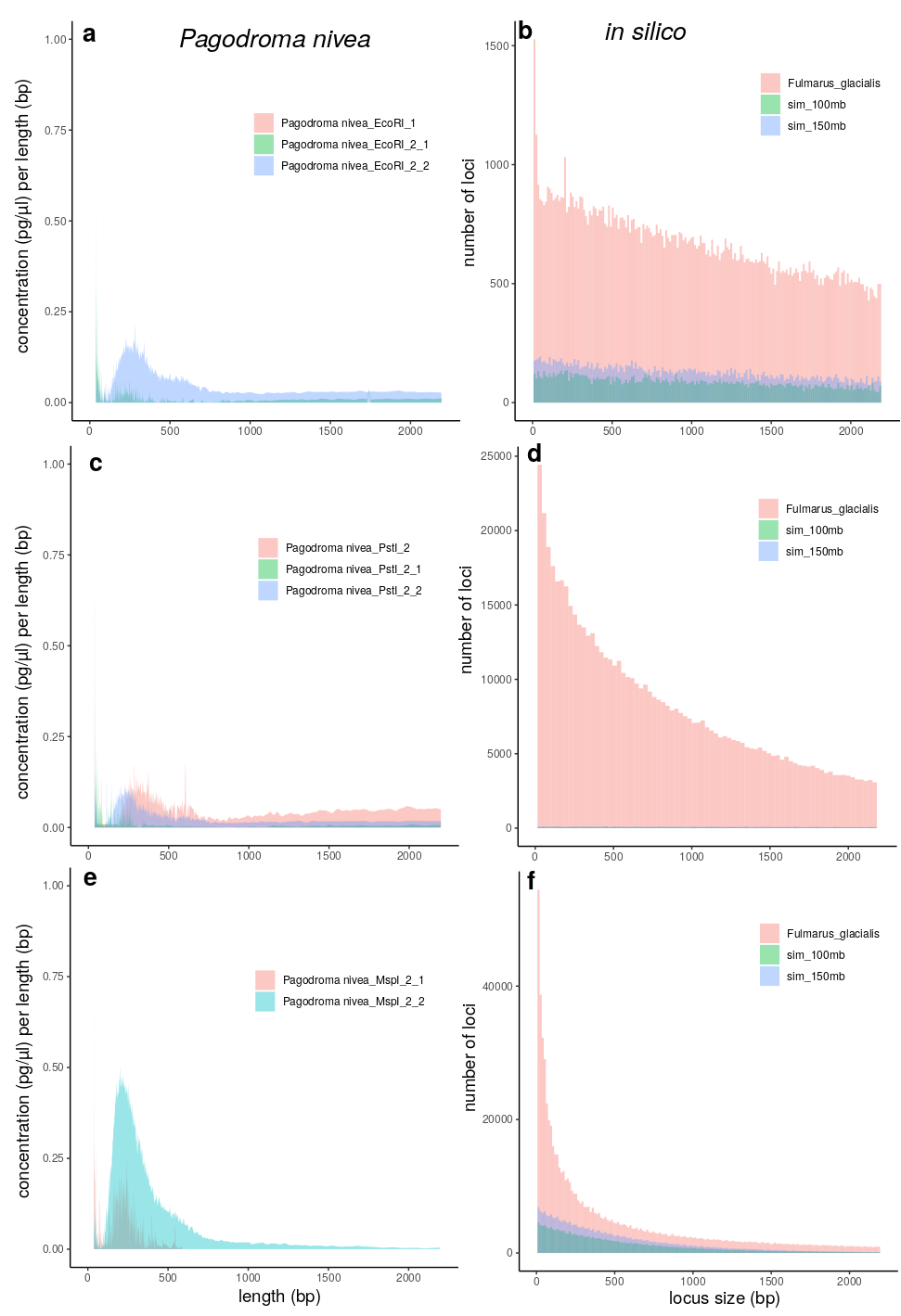

**Figure S3.5.** Empirical Bioanalyzer results with digested DNA are shown as concentration over fragment size (a, c, e) and estimated loci numbers over locus size from *in silico* digestions (b, d, f). The tests were conducted with restrictions enzymes *EcoRI* (a, b), *PstI* (c, d) and *MspI* (e, f). Results for the bird species *Pagodroma nivea* are shown next to *in silico* estimates using a related reference genome of *Fulmarus glacialis* and two simulated genomes of 100 and 150 Mb size (note that this was the absolute size used for *in silico* computations, but resulting estimates were extrapolated to 1000 and 1500 Mb).

**Additional File 4. DOCX. Reduced representation sequencing (RRS) setups for seven individually optimized protocols.** These setups were optimized in order to be run on a HiSeq 4000 platform (Illumina). The choice of restriction enzyme(s) and size window was optimized to obtain approximately 30× coverage (or half that in a worst-case scenario) with the assumed genome size (conservatively estimated based on available information, see Table 1). Marker density was estimated as a comparable measure to the metastudy by Lowry et al. (40).

| Class | Target Species | Restriction Enzyme (Combination) | Size Window (bp) | Assumed Genome Size (Mb) | Coverage^†^ | Marker Density^†^ (bp per 1 SNP) |
| --- | --- | --- | --- | --- | --- | --- |
| *Ostracoda* | Macrocyprididae | *ApeKI* | 250-500 | 250 | 35.3× | 941 |
| *Malacostraca* | *Charcotia obesa* | *SbfI_MspI* | 250-450 | 27,000 | 34.0× | 97,904 |
|  | *Eusirus pontomedon* | *EcoRI_SphI* | 250-350 | 7,000 | 30.7× | 22,898 |
| *Bivalvia* | *Laternula elliptica* and *Aequiyoldia eightsii* | *ApeKI* | 250-350 | 3,000 | 29.7 – 37.4× | 9,492 – 11,965 |
| *Asteroidea* | *Bathybiaster loripes* and *Psilaster charcoti* | *ApeKI* | 250-400 | 500 | 31.6 – 39.4× | 1,685 – 2,100 |
| *Actinopterygii* | *Trematomus bernacchii* and *T. loennbergii* | *EcoRI_MspI* | 250-600 | 1,500 | 33.3× | 4,944 |
| *Aves* | *Pagodroma nivea nivea* and *P. nivea confusa* | *PstI* | 250-400 | 1,500 | 33.8× | 5,410 |

^†^ assuming 300 million reads of 150 bp length spread over 96 individuals and 0.01 SNP/bp

**Additional File 5. Reduced representation sequencing (RRS) laboratory protocol based on the protocol from Peterson et al. (17).** The protocol is scaled for use with 192 samples and with restriction enzymes *EcoRI* and *MspI*; the reagent volumes can be scaled down/up to suit other sample numbers; if other enzymes are used, the respective reaction conditions must be adjusted.

**Step 1.** Prepare two PCR plates with 15 µL of each DNA sample at a concentration of 20 ng/µL

**Step 2.** Restriction enzyme digestion of 192 samples with *EcoRI*-HF and *MspI* (NEB, New England Biolabs)

- Prepare master mix for 220 samples:
  - Cut Smart buffer (NEB, 2 µL per sample): 440 µL
  - *EcoRI*-HF (1 µL per sample): 220 µL
  - *MspI* (1 µL per sample): 220 µL
  - Molecular grade water (1 µL per sample): 220 µL
- Vortex master mix, spin down briefly and put on ice
- Distribute 135 µL of the mix in 8 well strip and add 5 µL to each well of the sample plates with multipipet
- Total volume in each well: 20 µL
- Incubate for **3 h at 37° C**, cool down to 10° C
- Heat inactivation: **20 min at 65° C**, cool down to 10° C

**Step 3.** Ligation (T4 DNA ligase from NEB)

- Prepare master mix for 220 samples in falcon tube
  - Cut Smart buffer (NEB, 2 µl per sample): 440 μL
  - T4 ligase (1 µl per sample): 220 μL
  - *MspI* adaptor (9 µM; 2 µl per sample): 440 µl
  - rATP (1 µl per sample): 220 µl
  - Molecular grade water (12 µl per sample): 2640 μL
- Distribute in a clean plastic tray and add 18 μL to each well containing digested DNA with multipipet
- Total volume in each well: 38 μL
- Add 2 µl *EcoRI* adaptor (0.6 µM) to each sample: **watch out, there are 8 different adaptors with 8 barcodes**. Put adaptor 1 in wells A1, A2, …till A12. Adaptor 2 in wells B1, B2, … till B12 and the same for the others
- Incubate **30 min at 22° C**, followed by **10 min at 65°C**

**Step 4.** Purification with CleanPCR beads (CleanNA; GC Biotech);

to reduce costs of CleanPCR beads, only 20 μL will be purified

- Add 20 μL beads to new plate
- Add 20 μL of digestion/ligation mixture
- Mix by carefully pipetting up and down 10 times to ensure proper mixing
- Incubate 5 min at room temperature
- Place plate on magnet for 5 min to separate beads from solution
- Remove 35 μL of the clear solution while the plate is still on the magnet. Discard solution. Avoid taking out any beads; leave ca. 5 μL of the solution behind.
- Add 200 μL of 70% ethanol and wait 30 s
- Remove 200 μL ethanol (beads are now attached much better to the wall)
- Add 200 μL of 70% ethanol, wait 30 s
- Remove all supernatant (230 μL of ethanol). Check whether all ethanol is removed. Take 10 μL multipipet to double check whether all wells are empty. Residual ethanol may interfere with downstream PCR
- Remove plate from magnet and add 40 μL elution buffer (e.g. from Qiagen kit) or pure water (Sigma)
- Mix by pipetting 10 times up and down
- Incubate 5 min
- Put plate on magnet for 5 min to separate beads from solution
- Transfer 30 μL to new plate (make sure to not transfer beads, although they are not necessarily problematic later on)

**Step 5.** PCR on individual samples

- Prepare master mix for 200 samples
  - NEB Q5 hotstart master mix (12.5 µl per sample): 2500 µl
  - Molecular grade water (8.5 µl per sample): 1700 µl
  - F-Primer (5 µM, 1 µl per sample): 200 µl
- Distribute 22 μL of the mix and add 1 μL of R-primer (5 µM): **watch out, there are 12 different primers with 12 barcodes**. Put primer 1 in wells A1, B1, … till H1. Primer 2 in wells A2, B2, … till H2 and the same for the others.
- Add 2 µl of the purified digestion-ligation mix
- Total volume: 25 μL
- Initial denaturation at **98° C for 30 s** followed by 13 cycles of **10 s at 98° C**, **30 s at 65° C** and **30 s at 72°C**. Final elongation **5 min at 72°C**.

**Step 6.** Purification with CleanPCR beads

- Purify PCR product as in step 3 but add only 20 µL of beads to 25 µL PCR product (0.8 ratio)
- Follow protocol in step 3
- The final elution volume is 25 µL and 20 µL is transferred to a new tube

**Step 7.** Quantification

- Use the Quant-iT PicoGreen protocol (Thermo Fisher Scientific Inc.) and a microplate reader or a similar photometric method to precisely quantify the individual, amplified ddRAD samples following the manufacturer’s instructions.
- If DNA quantity at this step is too low, one may try to go back to step 5 and try with more PCR cyles. This also increases the amount of PCR duplicates of course.

**Step 8.** Pooling of the samples

- Depending on the lowest concentration take 20 ng (if possible, otherwise 10 ng or 5 ng) from each sample and transfer into one single tube.
- Quantify the pooled sample with PicoGreen again and check on gel.

**Step 9.** Export to KU Leuven Genomics Core.

The library/libraries are size selected (do not forget to add adaptor length to the chosen size window) and quantified at the Genomics Core using a Pippin Prep (Sage Science) and qPCR, respectively.

See original protocol version for further details:

Peterson, B.K., Weber, J.N., Kay, E.H., Fisher, H.S., Hoekstra, H.E. (2012) Double digest RADseq: an inexpensive method for *de novo* SNP discovery and genotyping in model and non-model species. PLoS One 7(5), e37135. <https://doi.org/10.1371/journal.pone.0037135>

**Additional File 6. DOCX. Reduced representation sequencing (RRS) laboratory protocol based on the protocol from Elshire et al. (20).** The protocol is scaled for use with 192 samples and with restriction enzymes *PstI* or *ApeKI*; the reagent volumes can be scaled down/up to suit other sample numbers; if other enzymes are used, the respective reaction conditions must be adjusted.

**Step 1.** Prepare two PCR plates with 10 µL of each DNA sample at a concentration of 10 ng/µL

**Step 2.** Restriction enzyme digestion of 192 samples with *PstI* or *ApeKI* (NEB, New England Biolabs)

- Add 6 μL of adaptor (1/10 diluted) to each well with multipipet
- Prepare master mix for 220 samples
  - NEB buffer 3 (2 μL per sample): 440 μL
  - *PstI* or *ApeKI* (1 μL per sample): 220 μL
  - Molecular grade water (1 μL per sample): 220 μL
- Vortex master mix, spin down briefly and put on ice
- Distribute 110 μL of the mix in 8 well strip and add 4 μL to each well of the sample plates with multipipet
- Total volume in each well: 20 μL
- For *PstI*: incubate **2 h at 37° C**, cool down to 10° C
- For *ApeKI*: incubate **2 h at 75° C**, cool down to 10° C

**Step 3.** Ligation (T4 DNA ligase from NEB)

- Prepare master mix for 220 samples in falcon tube
  - 10x T4 DNA ligase buffer (5 μL per sample): 1100 μL
  - T4 ligase (1.2 μL per sample): 264 μL
  - Molecular grade water (23.8 μL per sample): 5236 μL
- Distribute in a clean plastic tray and add 30 μL to each well containing digested DNA with multipipet
- Total volume in each well: 50 μL
- Incubate **1 h at 22° C**, followed by **30 min at 65° C** (heat inactivation of enzyme)

**Step 4.** Purification with CleanPCR beads (CleanNA; GC Biotech);

to reduce costs of CleanPCR beads, only 25 μL will be purified

- Add 25 μL beads to new plate
- Add 25 μL of digestion/ligation mixture
- Mix by carefully pipetting up and down 10 times to ensure proper mixing
- Incubate 5 min at room temperature
- Place plate on magnet for 5 min to separate beads from solution
- Remove 45 μL of the clear solution while the plate is still on the magnet. Discard solution. Avoid taking out any beads; leave ca. 5 μL of the solution behind.
- Add 200 μL of 70% ethanol and wait 30 s
- Remove 200 μL ethanol (beads are now attached much better to the wall)
- Add 200 μL of 70% ethanol, wait 30 s
- Remove all supernatant (230 μL of ethanol). Check whether all ethanol is removed. Take 10 μL multipipet to double check whether all wells are empty. Residual ethanol may interfere with downstream PCR
- Remove plate from magnet and add 40 μL elution buffer (e.g. from Qiagen kit) or pure water (Sigma)
- Mix by pipetting 10 times up and down
- Incubate 5 min
- Put plate on magnet for 5 min to separate beads from solution
- Transfer 35 μL to new plate (make sure to not transfer beads, although they are not necessarily problematic later on)

**Step 5**. PCR on separate samples

- Prepare master mix for 200 samples
  - NEB Q5 hotstart master mix (12.5 µL per sample): 2500 µL
  - Molecular grade water (10.5 µL per sample): 2100 µL
  - Primer mix (contains F and R primer, each at 5 µM, 1 µL per sample): 200 µL
- Distribute 24 μL of the mix and add 1 μL of cleaned ligation product
- Total volume: 25 μL
- Initial denaturation at **98° C for 30 s**, followed by 18 cycles of **10 s at 98° C, 30 s at 65° C** and **30 s at 72° C**. Final elongation **5 min at 72° C**.

**Step 6.** Purification with CleanPCR beads

- Purify PCR product as in step 3 but add only 20 µL of beads to 25 µL PCR product (0.8 ratio)
- Follow protocol in step 3
- The final elution volume is 30 µL and 25 µL is transferred to a new tube

**Step 7.** Quantification

- Use the Quant-iT PicoGreen protocol (Thermo Fisher Scientific Inc.) and a microplate reader or a similar photometric method to precisely quantify the individual, amplified ddRAD samples following the manufacturer’s instructions.
- If DNA quantity at this step is too low, one may try to go back to step 5 and try with more PCR cyles. This also increases the amount of PCR duplicates of course.

**Step 8.** Pooling of the samples

- Depending on the lowest concentration take 5 or 10 ng from each sample and transfer into one single tube.
- Quantify the pooled sample with PicoGreen again and check on gel.

**Step 9.** Export to KU Leuven Genomics Core.

The library/libraries are size selected (do not forget to add adaptor length to the chosen size window) and quantified at the Genomics Core using a Pippin Prep (Sage Science) and qPCR, respectively.

See original protocol version for further details:

Elshire, R.J., Glaubitz, J.C., Sun, Q., Poland, J.A., Kawamoto, K., Buckler, E.S., Mitchell, S.E. (2011) A robust, simple genotyping-by-sequencing (GBS) approach for high diversity species. PLoS One 6(5), e19379. <https://doi.org/10.1371/journal.pone.0019379>

**Additional File 7. DOCX. Results from parameter optimization for *de novo* assembly and genotyping.** Eight parameter optimization series were conducted following Rochette & Catchen (42) to identify optimal parameters to genotype reduced representation sequencing (RRS) data with Stacks v2.4 (15); one test series for each species/species complex. The Stacks parameter m was kept constant (m = 3), while parameters M and n were varied together from 1 to 9. Subsequently, only loci present in 80 % of the samples were retained and for each M=n parameter the number of loci and polymorphic loci was plotted, as well as the proportion of these loci containing 0 to 10 or >10 SNPs. In ostracods, the library contained DNA from a species-complex, resulting in very few shared loci across 80 % of the samples. Therefore, in this case results based on loci shared by 50 % of samples are shown. Optimal M=n values were decided in all cases with this information (and reported in Table 4). Note, however, that it is impossible to make absolute calls regarding the ideal value.

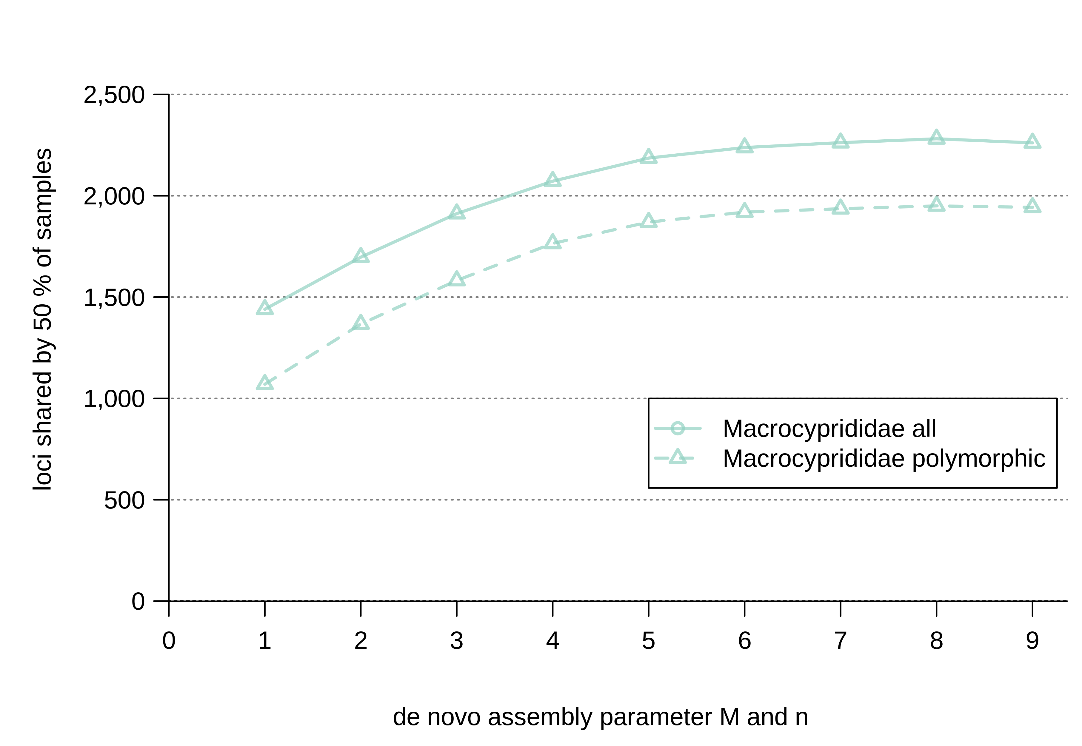

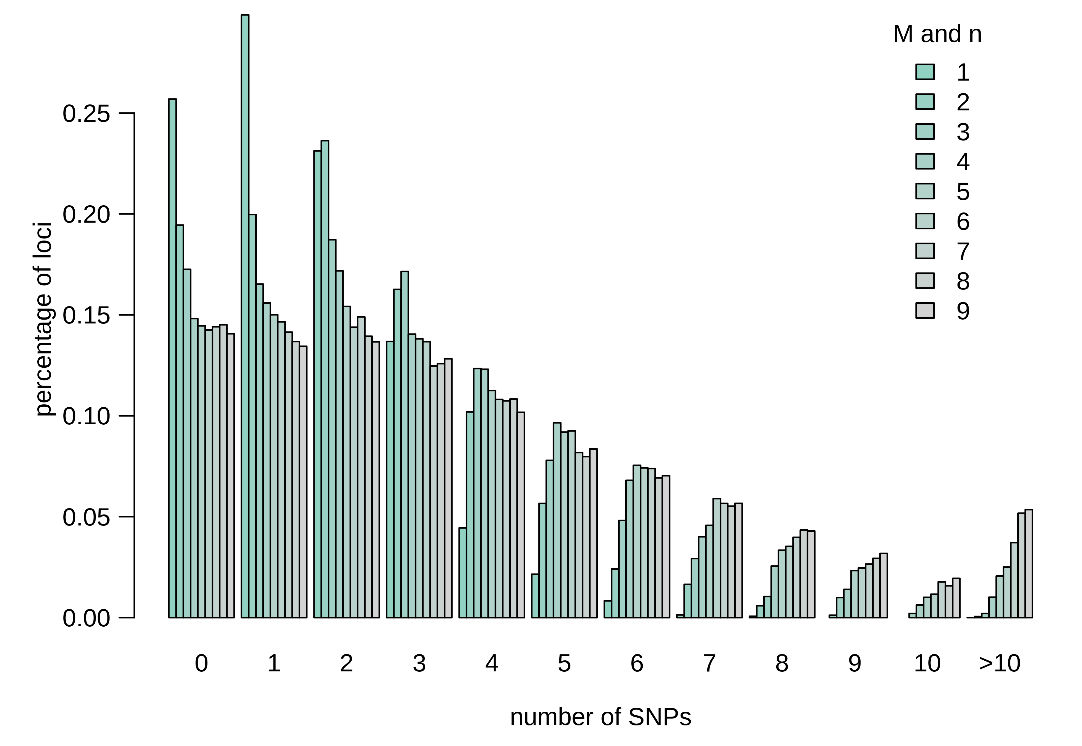

**Figure S7.1.** Number of loci and polymorphic loci shared by 50 % of samples from test library 1 across nine values for parameter M and n in Stacks v.2.4 (top) and number of SNPs per locus across the same parameter range (bottom). M = n = 6 was retained. Note that library 1 contained a species complex (*Macroscapha opaca-tensa* complex) with only few loci shared across many samples. Therefore, results of loci shared by only 50 % of the samples are shown.

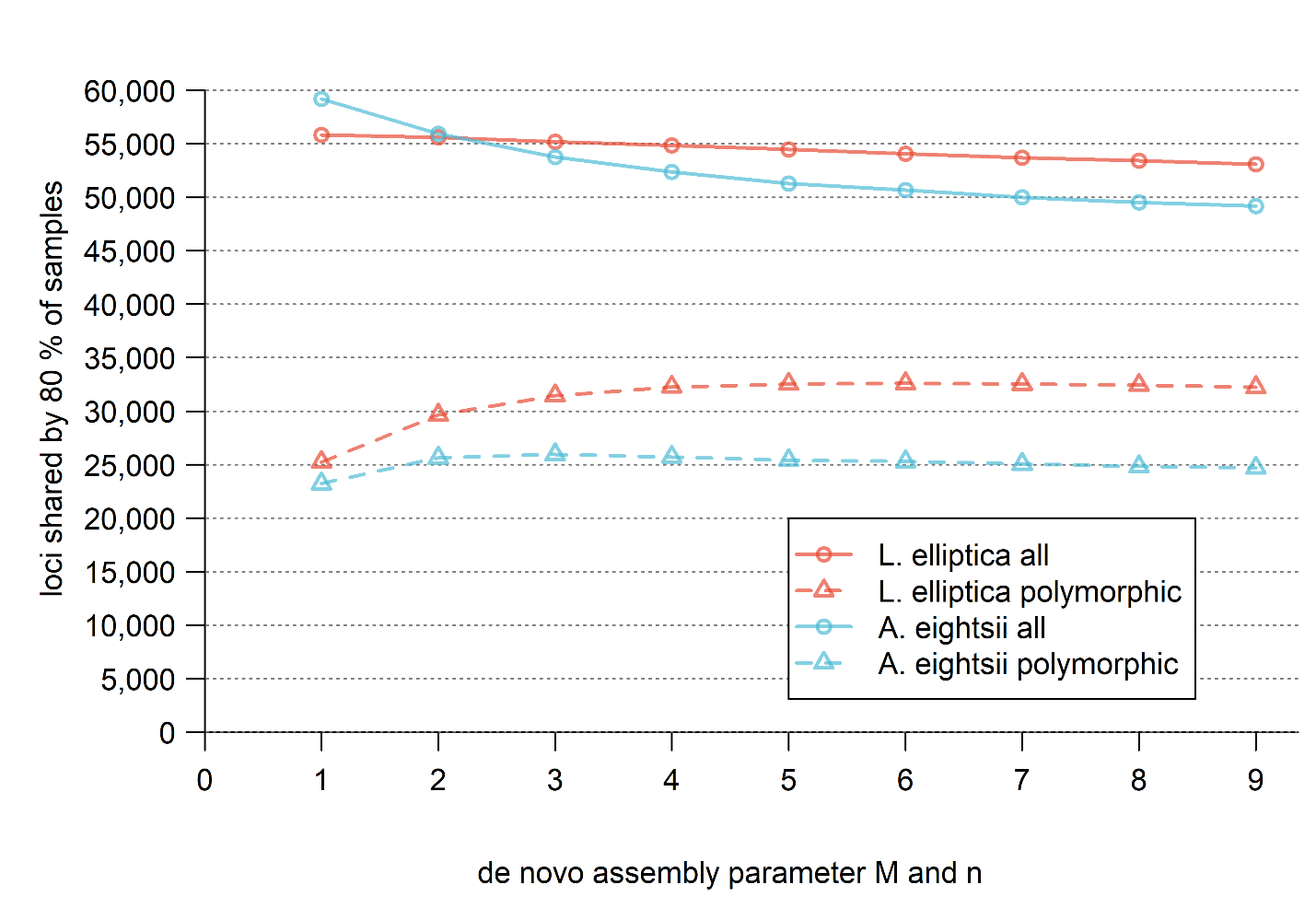

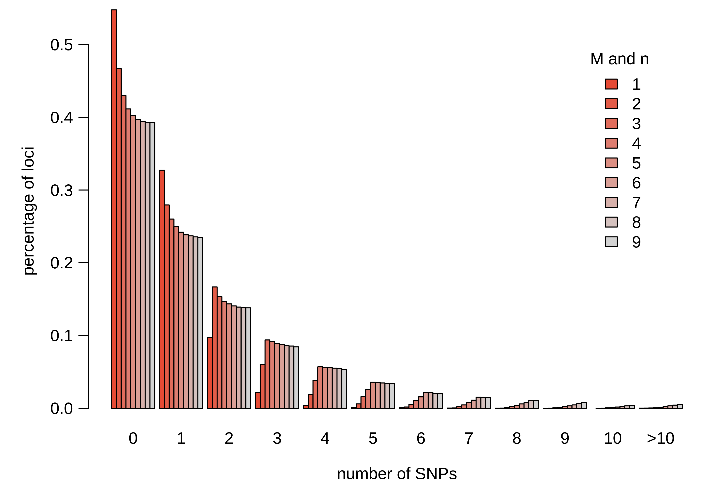

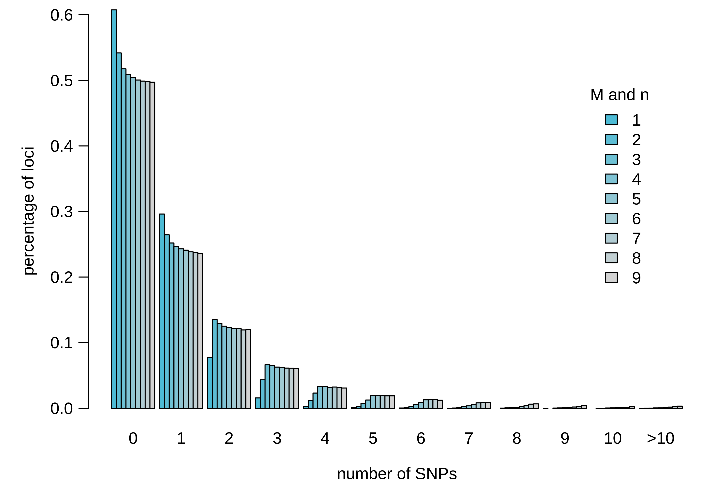

**Figure S7.2.** Number of loci and polymorphic loci shared by 80 % of samples from test library 2 across nine values for parameter M and n in Stacks v.2.4 (top) and number of SNPs per locus across the same parameter range for *Laternula elliptica* (bottom left) and *Aequiyoldia eightsii* (bottom right). M = n = 4 was retained.

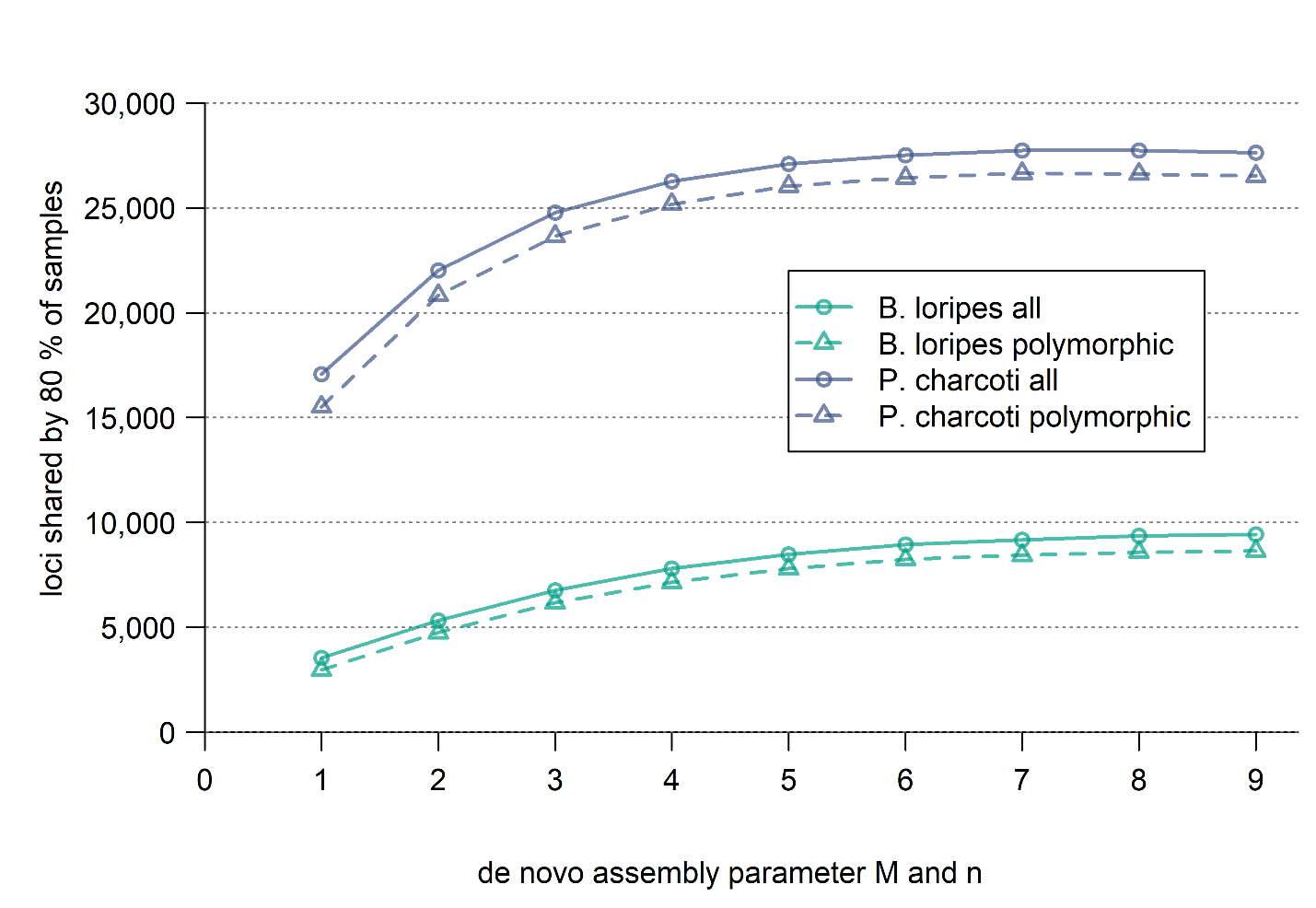

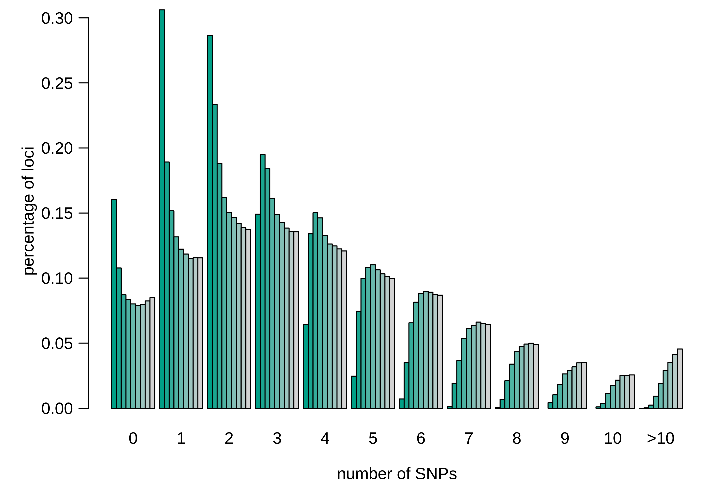

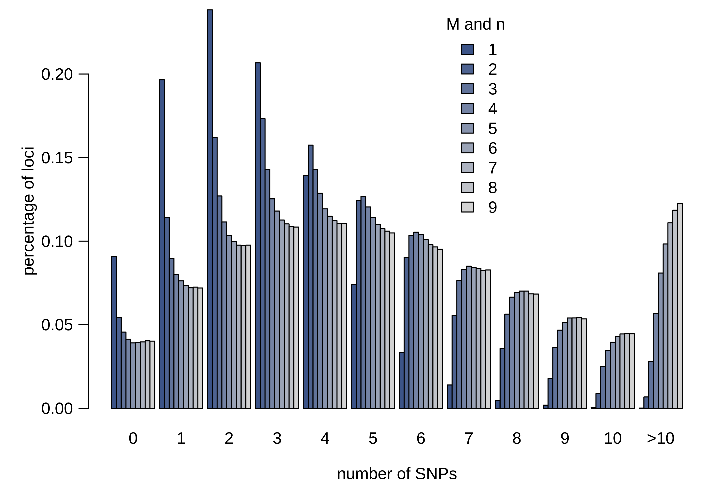

**Figure S7.3.** Number of loci and polymorphic loci shared by 80 % of samples from test library 3 across nine values for parameter M and n in Stacks v.2.4 (top) and number of SNPs per locus across the same parameter range for *Bathybiaster loripes* (bottom left) and *Psilaster charcoti* (bottom right). M = n = 5 was retained.

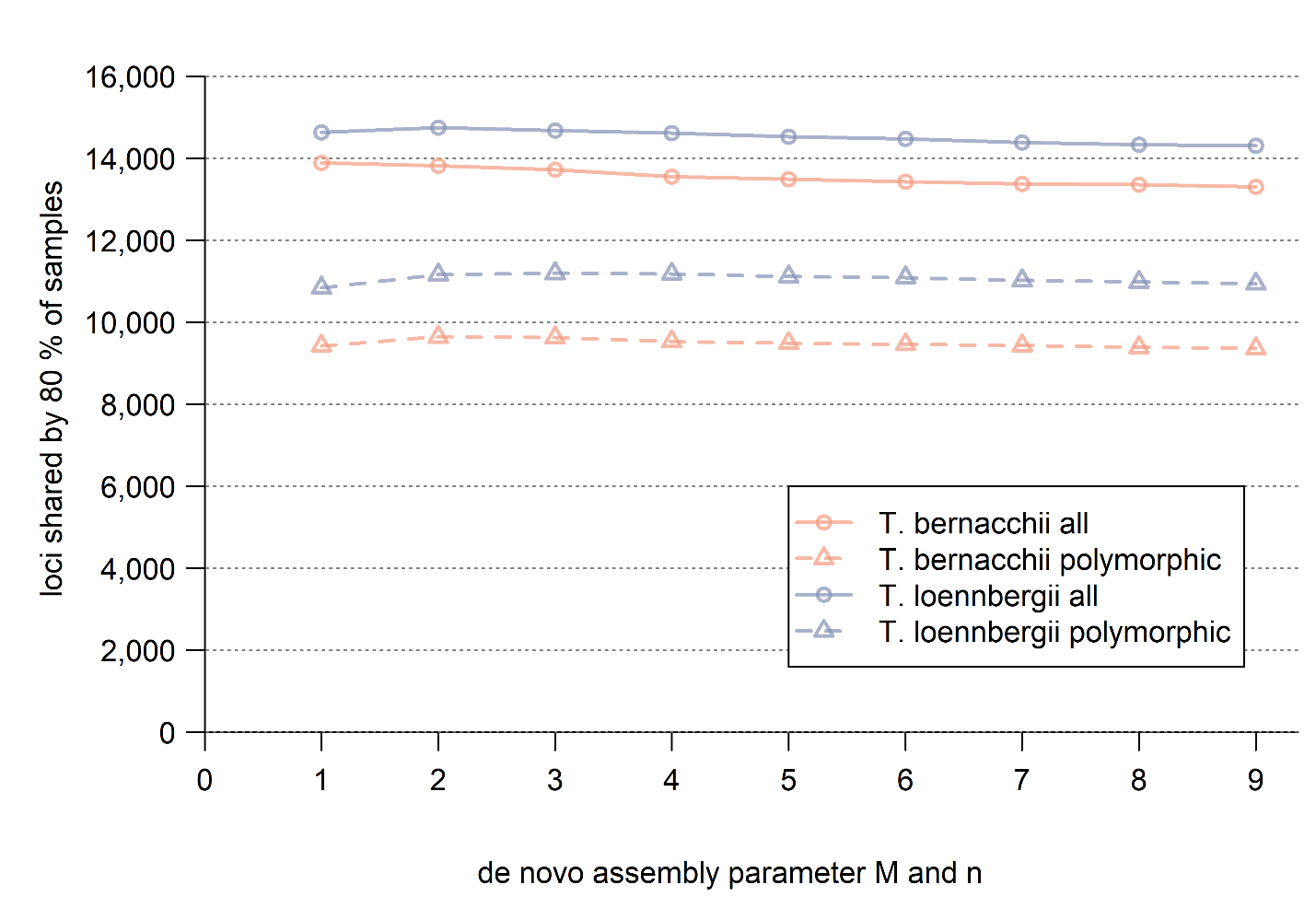

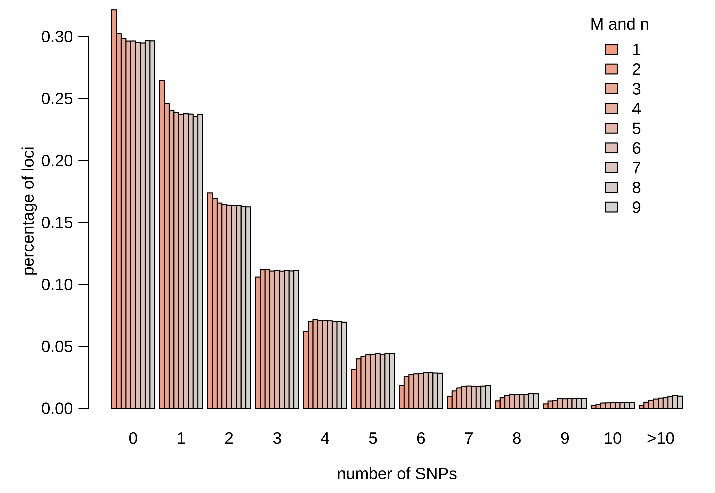

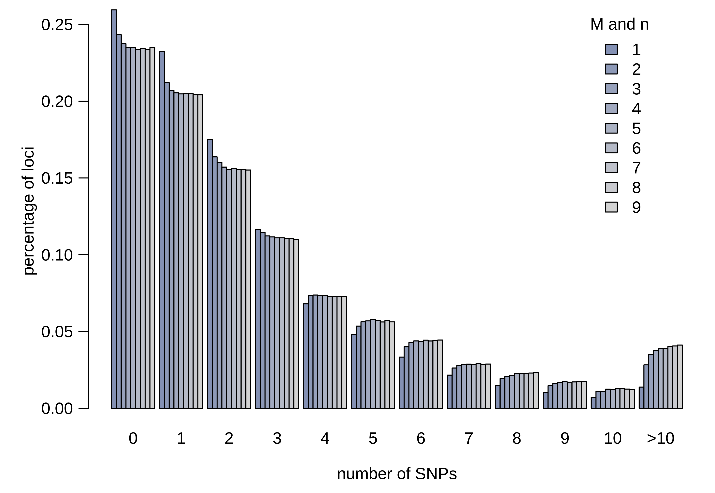

**Figure S7.4.** Number of loci and polymorphic loci shared by 80 % of samples from test library 4 across nine values for parameter M and n in Stacks v.2.4 (top) and number of SNPs per locus across the same parameter range for *Trematomus bernacchii* (bottom left) and *T. loennbergii* (bottom right). M = n = 3 was retained.

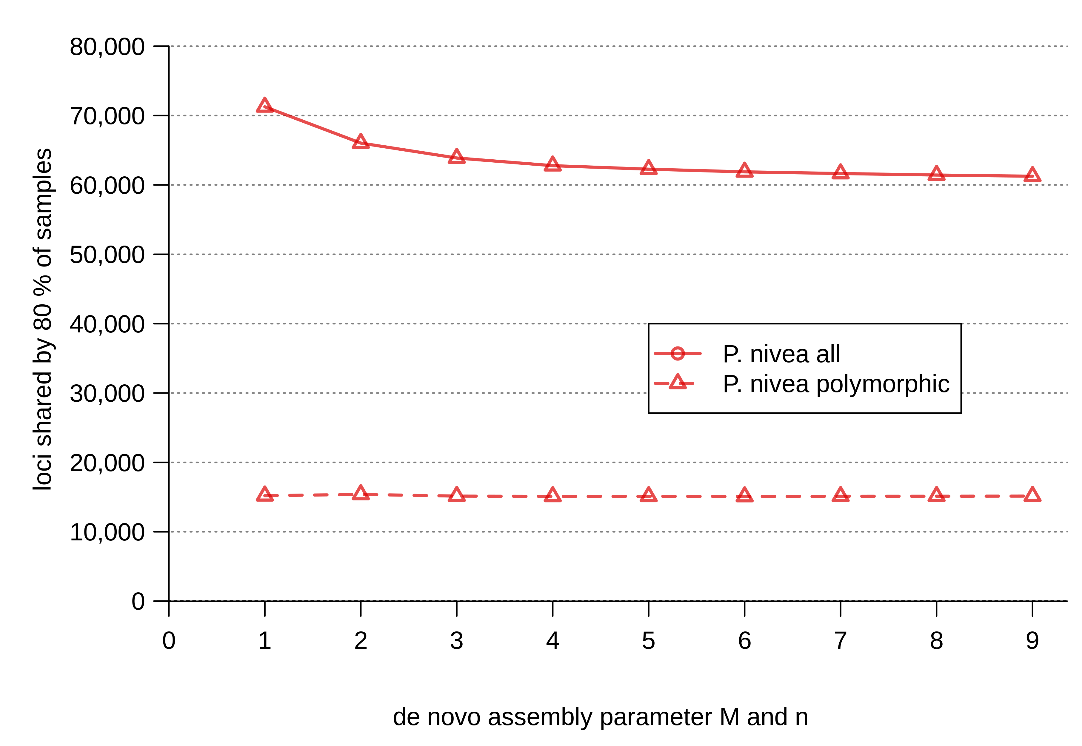

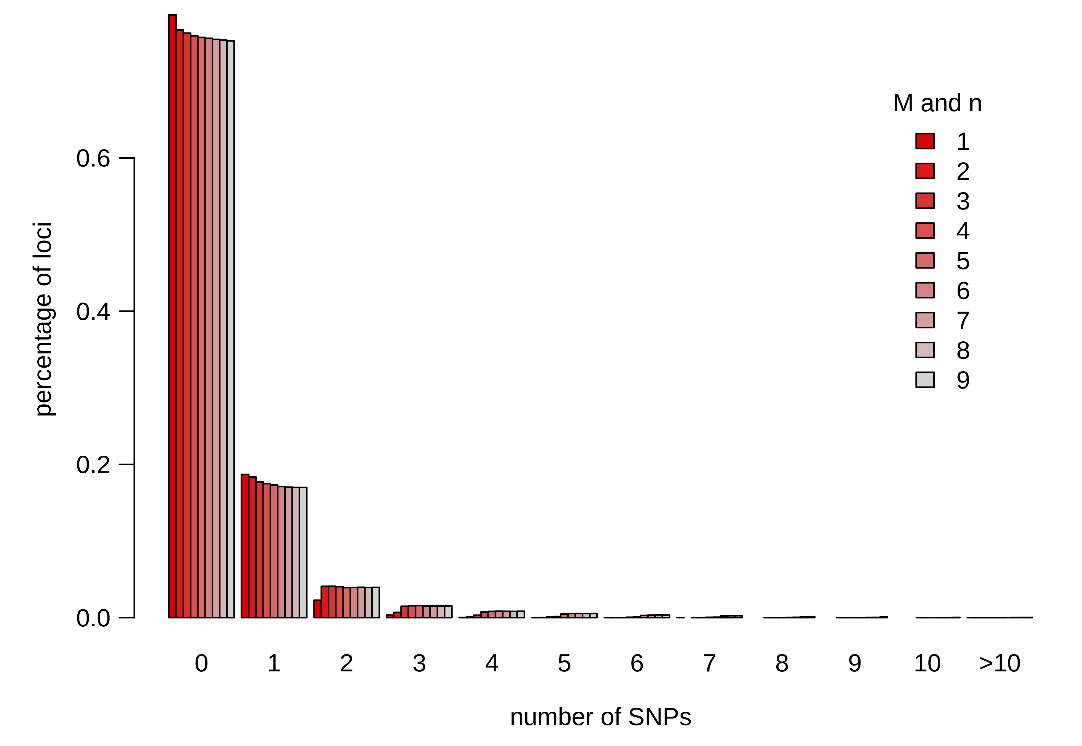

**Figure S7.5.** Number of loci and polymorphic loci shared by 50 % of samples from test library 5 across nine values for parameter M and n in Stacks v.2.4 (top) and number of SNPs per locus across the same parameter range (bottom). M = n = 3 was retained. Note that library 2 contained only few samples with likely high levels of degradation, possibly explaining the low amount of polymorphism detected.
